# LRRC8 complexes are adenosine nucleotide release channels regulating platelet activation and arterial thrombosis

**DOI:** 10.1101/2024.09.26.615233

**Authors:** John D. Tranter, Ryan T. Mikami, Ashutosh Kumar, Gavriel Brown, Tarek M. Abd El-Aziz, Yonghui Zhao, Nihil Abraham, Chloe Meyer, Abigail Ajanel, Litao Xie, Katrina Ashworth, Juan Hong, Haixia Zhang, Tripti Kumari, Adam Balutowski, Alice Liu, David Bark, Vinayak K. Nair, Nina M. Lasky, Yongmei Feng, Nathan O. Stitziel, Daniel J. Lerner, Robert A. Campbell, Jorge Di Paola, Jaehyung Cho, Rajan Sah

**Author notes:** Correspondence: Rajan Sah M.D. Ph.D. 425 S Euclid Blvd BJCIH 9609 St. Louis, MO 63105 Jaehyung Cho, Ph.D. 660 S. Euclid Ave, St. Louis, MO 63110. Equal Contribution Authors (John D. Tranter, Ryan T. Mikami, Ashutosh Kumar, Gavriel Brown).

## Abstract

Platelet shape and volume changes are early mechanical events contributing to platelet activation and thrombosis. Here, we identify single-nucleotide polymorphisms in Leucine-Rich Repeat Containing 8 (LRRC8) protein subunits that form the Volume-Regulated Anion Channel (VRAC) which are independently associated with altered mean platelet volume. LRRC8A is required for functional VRAC in megakaryocytes (MKs) and regulates platelet volume, adhesion, and agonist-stimulated activation, aggregation, ATP secretion and calcium mobilization. MK-specific LRRC8A cKO mice have reduced arteriolar thrombus formation and prolonged arterial thrombosis without affecting bleeding times. Mechanistically, platelet LRRC8A mediates swell-induced ATP/ADP release to amplify agonist-stimulated calcium and PI3K-AKT signaling via P2X1, P2Y_1_ and P2Y_12_ receptors. Small-molecule LRRC8 channel inhibitors recapitulate defects observed in LRRC8A-null platelets *in vitro* and *in vivo*. These studies identify the mechanoresponsive LRRC8 channel complex as an ATP/ADP release channel in platelets which regulates platelet function and thrombosis, providing a proof-of-concept for a novel anti-thrombotic drug target.

## Introduction

Platelets are essential for hemostasis and thrombosis ^1–3^. The process of thrombus formation involves three distinct stages: platelet adhesion, activation, and aggregation ^4^. Following vascular injury, platelets adhere to collagen and von Willebrand factor (vWF) through glycoprotein VI (GPVI) and the GPIb/IX/V complex, respectively. Platelets undergo rapid shape changes through cytoskeletal rearrangements ^5^ and release granule molecules such as ADP, serotonin, and thromboxane A2, which act on their respective G-protein coupled receptors (GPCRs) in autocrine and paracrine manners to promote thrombus formation ^6–8^. Whilst platelet shape change occurs early in the activation process and is considered to be required for platelet aggregation ^5^, the molecular mechanisms responsible for sensing and transducing these changes in platelet volume remain to be elucidated.

Cytosolic calcium levels contribute to platelet activation and are increased by calcium release from intracellular stores such as the dense tubular system as well as calcium influx across the plasma membrane ^9,10^. Calcium release from the dense tubular system occurs downstream of phospholipase C (PLC), which can be activated by interactions between ligands and their receptors. For example, ADP can activate P2Y_1_ and P2Y_12_ receptors resulting in calcium release and PI3K-AKT signaling, respectively, and activating αIIbβ3 integrin ^11,12^. A major pathway of calcium entry across the plasma membrane is that mediated by both STIM1-Orai1^13,14^ and P2X1^15^. P2X1 is an ATP-gated, calcium-permeant ion channel which, when activated by ATP released from damaged endothelium and/or activated platelets, contributes to increases in intracellular calcium to amplify platelet activation ^16,17^.

While ion channels are traditionally associated with excitable cell types, they are ubiquitous proteins which mediate ionic currents in almost all cells, including platelets. Various types of ion channels are implicated in platelet function, including ligand-gated cation channels, voltage and calcium-gated potassium channels, transient receptor potential (TRP) channels ^18–20^, and TMEM16F channels. TMEM16F channels are lipid scramblases which, when deficient, are responsible for Scott syndrome, a rare bleeding disorder ^21,22^. In addition, mechanosensitive and mechanoresponsive ion channels, such as Piezo1, have been recently described to regulate platelet function ^23^. Mechanotransduction is a process in which mechanical forces applied to cells results in translation into electrical signaling via the permeation of ions across the lipid bilayer and is mediated by mechanosensitive or mechanoresponsive ion channels, directly or indirectly gated by mechanical forces, respectively ^24^. LRRC8A (leucine-rich repeat-containing protein 8A, also known as SWELL1) is the essential subunit of the heterohexameric LRRC8 volume-regulated anion channel (VRAC) ^25,26^, which contain at least one LRRC8A subunit in combination with, B, C, D and/or E subunits. LRRC8 channels are mechanoresponsive anion channels expressed in many cell types, including adipocytes ^27^, endothelial cells ^28^, pancreatic β-cells ^29–31^, skeletal muscle cells ^32,33^, immune cells ^34–36^, astrocytes ^37–40^, microglia ^41–43^, and neurons ^44–47^. As LRRC8 channels are activated by cell swelling and shape change, we hypothesized that LRRC8s are expressed and functional in platelets and activate in response to platelet shape change at sites of vascular injury, contributing to arterial thrombosis. In the current study, we first identify a genetic signal for LRRC8 proteins as important regulators of platelet volume and function in humans. We show that LRRC8A-D proteins are expressed in murine and human platelets, form functional ATP and ADP-permeant VRAC currents in megakaryocytes (MKs), and contribute to platelet activation, adhesion, and aggregation, as well as arterial thrombosis *in vivo*. Mechanistically, the LRRC8 complex mediates ATP and ADP release to amplify agonist-stimulated calcium influx and PI3K-AKT signaling via P2X1, P2Y_1_ and P2Y_12_ receptors, respectively. Additionally, we highlight the potential opportunity for targeting LRRC8 channels by small-molecule LRRC8 inhibitors as a novel anti-thrombotic strategy, and demonstrate that dicoumarol, a historically used anticoagulant drug and precursor to warfarin, has direct anti-platelet effects by targeting LRRC8 channels.

## Results

### Genome-wide Association Studies (GWAS) identified three SNPs assigned to LRRC8 genes that are associated with mean platelet volume

Mean platelet volume (MPV) is a measured trait in many GWAS studies because it is calculated by most automated hematologic analyzers. Human LRRC8A-E are encoded by five genes on three chromosomes: LRRC8A on human chromosome 9q34.11, LRRC8B, C, and D in tandem on chromosome 1p22.2, and LRRC8E on chromosome 19p13.2 ^48^. We searched the National Health Genome Research Institute - European Bioinformatics Institute (NHGRI-EBI) catalog of genome-wide association studies (ebi.ac.uk/gwas/home) for SNPs and identified a number of significant associations between MPV and SNP variants assigned to LRRC8 genes, including three listed in **Table 1**. Compared to the ancestral allele, the SNP variants assigned to LRRC8A and C are associated with increased MPV, and the variant assigned to D is associated with decreased MPV.

**Table 1.**
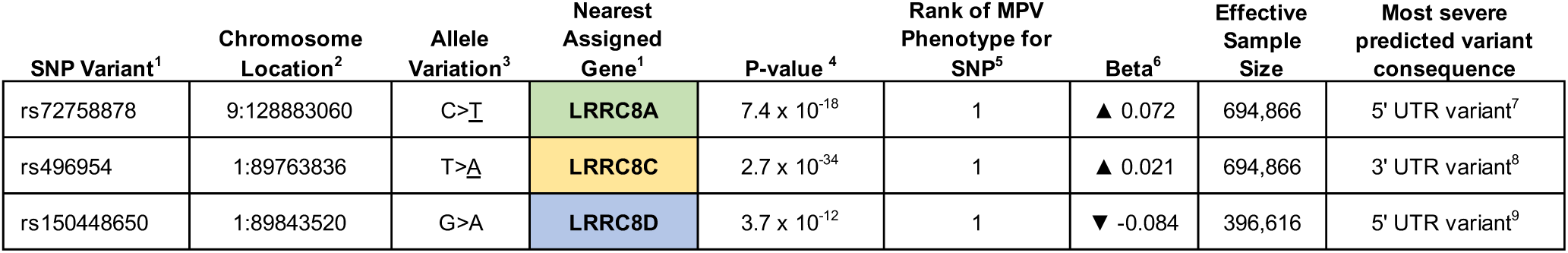
Single nucleotide polymorphisms in genes encoding LRRC8 proteins are associated with mean platelet volume. ^1^ SNP variants associated with mean platelet volume in the NHGRI-EBI Catalog of human genome-wide association studies (GWAS CAT; ebi.ac.uk/gwas/home). ^2^ Nucleotide position in the Genome Reference Consortium human build 38 (GRCh38) (chromosome: nucleotide; ncbi.nlm.nih.gov/snp/) ^3^ Ancestral allele > variant allele (ncbi.nlm.nih.gov/snp/) ^4^ *p* value for the association between the SNP variant and mean platelet volume phenotype (gwas.mrcieu.ac.uk/phewas/) ^5^ Rank of the association between the SNP variant and mean platelet volume phenotype, from most (1) to least significant (gwas.mrcieu.ac.uk/phewas/) ^6^ Per unit change in mean platelet volume associated with the SNP variant ^7^ ncbi.nlm.nih.gov/snp/?term=rs72758878 ^8^useast.ensembl.org/Homo_sapiens/Variation/Explore?db=core;r=1:89763336-89764336;v=rs496954;vdb=variation;vf=25209 ^9^useast.ensembl.org/Homo_sapiens/Variation/Explore?r=1:89843020-89844020;v=rs150448650;vdb=variation;vf=2698779

### Phenome-wide Association Studies (PheWAS) identified mean platelet volume as the phenotype most closely associated with all three SNPs assigned to LRRC8 genes

To determine the relative strength of the association of the MPV phenotype with each variant, we conducted a PheWAS search for phenotypes associated with each SNP variant and found that MPV was the phenotype most closely associated with all three variants, with *p* values ranging from 3.7 x 10^-12^ to 2.7 x 10^-34^ (**Table 1**).

Because LRRC8 channels are heterohexamers comprised of different combinations of five LRRC8 subunits, and MPV is the most closely associated phenotype with exonic SNPs in genes assigned to three of these subunits located on two different chromosomes, we interpret these human genetic data as evidence that LRRC8 channels are involved in regulating platelet volume and function.

### LRRC8A-D transcripts are expressed in platelets and are associated with altered agonist-induced aggregation

To determine if LRRC8A-E transcripts are expressed in human platelets, we searched a publicly available plateletomics database (plateletomics.com, utilizing data from Rowley *et al.* ^49^ and Simon *et al.* ^50^) which details both human and murine platelet transcriptomics. *LRRC8A*, *LRRC8B*, *LRRC8C*, and *LRRC8D* transcripts are all expressed in platelets (top 75^th^, 13^th^, 42^nd^, and 5^th^ percentile of 5,911 transcripts, respectively) (**Supplementary** Fig. 1A). *LRRC8E* is not detected. Absolute LRRC8 transcript expression levels in human platelets reveal *LRRC8D*>*LRRC8B*>*LRRC8C*>*LRRC8A* (**Fig. 1A**), while in murine platelets *Lrrc8c*>*Lrrc8b*>*Lrrc8d*>*Lrrc8a* (**Fig. 1D**). In both human and mouse platelets, LRRC8 mRNA expression levels are substantially higher than mechanosensitive *PIEZO1*, *PIEZO2,* and other mechanoresponsive ion channels (*TRPV4*), and comparable to the highly expressed transmembrane protein *TMEM16F* (*Ano6*) that is important for platelet lipid scrambling and causal in Scott syndrome (**Fig. 1B and 1E**) ^22^. Consistent with human mRNA expression data, LRRC8A, B, C, and D proteins are expressed in both human (**Fig. 1C**) and murine (**Fig. 1F**) platelets, while LRRC8E is not.

**Figure 1.**
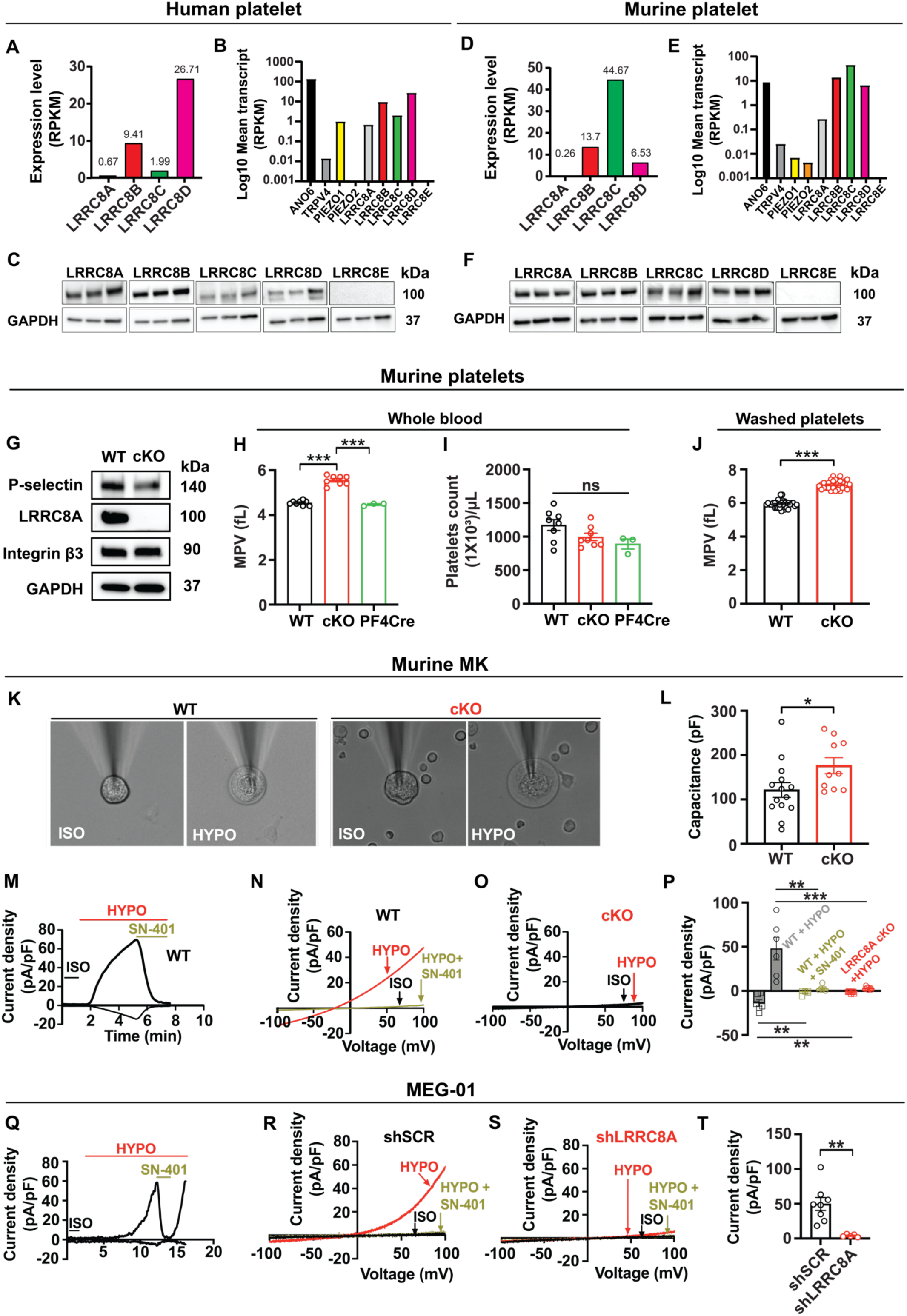
LRRC8 proteins are highly expressed in platelets, functionally encode VRAC in megakaryocytes and regulate platelet volume. **a**, **d** LRRC8 mRNA transcript expression in human (**a**) and mouse (**d**) platelets. **b**, **e** LRRC8 mRNA transcript expression compared to transcripts for ANO6, TRPV4, Piezo1 and Piezo2 in human (**b**) and mouse (**e**) platelets (**a – e** are data from Rowley *et al.* [49] - Supplemental Table 4). **c**, **f** Western blot showing LRRC8 subunit protein expression of human (**c**, n = 3) and WT C57BL/6N mouse (**f**, n = 3) platelets. **g** Western blot detecting P-selectin, LRRC8A, integrin β3, and GAPDH in platelets isolated from *Lrrc8a^fl/fl^*(WT) and *Pf4-Cre;Lrrc8a^fl/fl^* (cKO) mice. **h, i** Mean platelet volume (MPV) in whole blood from WT (n = 8), cKO (n = 8), and heterozygous controls (*Pf4-Cre;Lrrc8a^fl/+^,* n = 3) (**h**) with corresponding platelet counts (**i**). **j** MPVs of washed platelets isolated from WT (n = 35) and cKO (n = 31) mice. **k** Representative images of perforated patch-clamped freshly isolated megakaryocytes from WT or cKO mice, under isotonic (ISO) and hypotonic (HYPO) conditions. **l** Cell capacitances of megakaryocytes isolated from WT (n = 14) and cKO mice (n = 10). **m** VRAC current-time relationship of WT MK cell induced by hypotonic (210 mOsm) swelling, and subsequent inhibition upon application of 10 µM SN-401 (DCPIB). **n-o** VRAC current-voltage relationship after hypotonic swelling during voltage ramps from -100 mV to +100 mV in MKs isolated from WT +/- SN-401 (**n**) and cKO (**o**) mice. **p** Mean outward (+100 mV) and inward (-100 mV) current densities measured in WT MKs following hypotonic stimulation in the presence (*gold*: n = 6) or absence (*gray*: n = 6) of 10 µM SN-401, or in cKO MKs following hypotonic stimulation (*red:* n = 9). **q** VRAC current-time relationship in a MEG-01 cell induced by hypotonic swelling, and subsequent inhibition upon application of 10 µM SN-401. **r, s** VRAC current-voltage relationship during voltage ramps from -100 mV to +100 mV in MEG-01 cells transduced for 72h with either an adenoviral short hairpin control (shSCR; **r**) or LRRC8A (shLRRC8A; **s**) after hypotonic swelling in the presence or absence of 10 µM SN-401. **t** Mean current densities of shSCR (n = 8) and shLRRC8A (n = 5) transduced MEG-01 cells at +100 mV following hypotonic swelling. Data are represented as mean ± S.E.M. Statistical significance was determined by ordinary one-way ANOVA with Tukey’s multiple comparisons test for **h** and **i**, and by unpaired T-test for **j**, **l**, **p**, and **t**. ns – not significant; * *p* < 0.05; ** *p* < 0.01; *** *p* < 0.001.

Multiple regression analyses of human platelet aggregation in response to stimulation with adenosine diphosphate (ADP), protease-activated receptor 1 activating peptide (PAR1-AP), PAR4-AP, or arachidonic acid (AA), adjusting for mRNA level, age, gender, race, BMI, and platelet count identified a complex relationship between LRRC8 subunit mRNA levels and agonist-induced platelet aggregation. LRRC8A mRNA levels are significantly associated with enhanced ADP-and PAR4-AP-stimulated aggregation; LRRC8B mRNA levels are associated with reduced ADP-and PAR1-AP-stimulated aggregation; and LRRC8D mRNA levels are associated with reduced PAR1-AP-induced aggregation (**Supplementary** Fig. 1B). Taken together, these data provide correlative evidence to suggest a modulatory role for VRAC and LRRC8 proteins in regulating platelet function in humans.

### LRRC8A functionally encodes VRAC in MKs and regulates platelet volume

To explore the biological mechanisms implied by the human genetic studies, we generated MK-specific LRRC8A conditional knockout (cKO) mice by crossing *Lrrc8a*^fl/fl^ mice (*Swell1*^fl/fl^) ^27–29,32,51^ with platelet factor 4-Cre (*Pf4-Cre*) mice to generate *Lrrc8a^fl/fl;Pf4-cre+^* (cKO) and littermate control *Lrrc8a*^fl/fl^ mice (WT). LRRC8A protein is completely deleted in cKO mouse platelets while maintaining expression of integrin β3 and P-selectin (**Fig. 1G**). Consistent with the increase in MPV associated with the SNP assigned to LRRC8A in GWAS/PheWAS studies (**Table 1**), platelet-targeted LRRC8A deletion also increases MPV compared to both WT and platelet-targeted heterozygous controls (*Lrrc8a^fl/+;Pf4-cre+^*, **Fig. 1H&J**), without affecting platelet counts (**Fig. 1I**), or any other hematological parameters (**Supplementary** Fig. 2). Thus, LRRC8A deletion in mouse platelets increases in MPV, which phenocopies human genetic data (**Table 1**) implicating LRRC8 proteins as regulators of platelet volume.

To confirm that LRRC8 proteins form functional VRAC channels in platelets, we measured VRAC in freshly isolated murine MKs under isotonic conditions and with hypotonic swelling (**Fig. 1K**). The cell capacitance of LRRC8A cKO MKs (176 ± 17.9 pF, n = 10) are notably larger than WT MK (121 ± 16.6 pF, n = 14, *p* < 0.05) (**Fig. 1L**). WT MK develop robust VRAC in response to hypotonic swelling, and this is completely inhibited by 10 μM SN-401 (DCPIB) and abolished upon LRRC8A deletion (**Fig. 1M-P**), confirming this volume-sensitive, mechanoresponsive, SN-401-inhibited current to be LRRC8A-mediated. Similarly, robust VRAC currents are elicited in response to hypotonic swelling in the human megakaryoblast MEG-01 cell line and fully inhibited by 10 μM SN-401 (**Fig. 1Q&R**). Importantly, adenovirally-delivered shRNA-mediated LRRC8A knockdown (KD) (Ad-shLRRC8A-mCherry) significantly diminished MEG-01 VRAC (**Fig. 1S**), as compared to a scrambled control (Ad-shSCR-mCherry, **Fig. 1R&T**).

### LRRC8A is required for normal platelet adhesion, granule release, and aggregation

Next, we investigated whether LRRC8A regulates platelet function. Platelet adhesion is mediated by GPVI and the GPIb/IX/V complex that interact with collagen and vWF, respectively. Then, platelet aggregation is mediated by the interaction between activated αIIbβ3 integrin and fibrinogen. LRRC8A depletion markedly impairs platelet adhesion to fibrillar collagen-coated surface under shear conditions (650 s^−1^) (**Fig. 2A&B**, **Video S1&2**), and reduces thrombin-stimulated platelet activation as assessed by P-selectin exposure, a measure of α-granule release (**Fig. 2C**) and αIIbβ3 integrin activation (**Fig. 2D**), while activation with Ca^2+^ ionophore A23187 is preserved. To investigate the function of LRRC8A in human cells, we used CRISPR-edited human MKs generated from CD34^+^ cells (hiPSC MKs) isolated from cord blood ^52^ (**Fig. 2E**). Dual-guide RNAs (gRNAs) targeting LRRC8A in hiPSC MKs significantly depleted LRRC8A expression as compared to cells treated with non-targeting control gRNA (**Fig. 2F**). In LRRC8A KO hiPSC MKs, thrombin-stimulated P-selectin exposure is significantly reduced (**Fig. 2G**). Similarly, binding of PAC-1, a pentameric IgM which binds to activated αIIbβ3 integrin ^53^, is also significantly decreased in LRRC8A KO hiPSC MK cells (**Fig. 2H**). Platelet-specific LRRC8A depletion also impairs platelet aggregation in response to multiple agonists, including thrombin (**Fig. 2I**), ADP (**Fig. 2J**), collagen-related peptide (CRP, **Fig. 2K**) and a thromboxane A2 analogue (U46619, **Fig. 2L**). Similarly, ATP secretion is impaired in response to all agonists tested (**Fig. 2M-P**).

**Figure 2.**
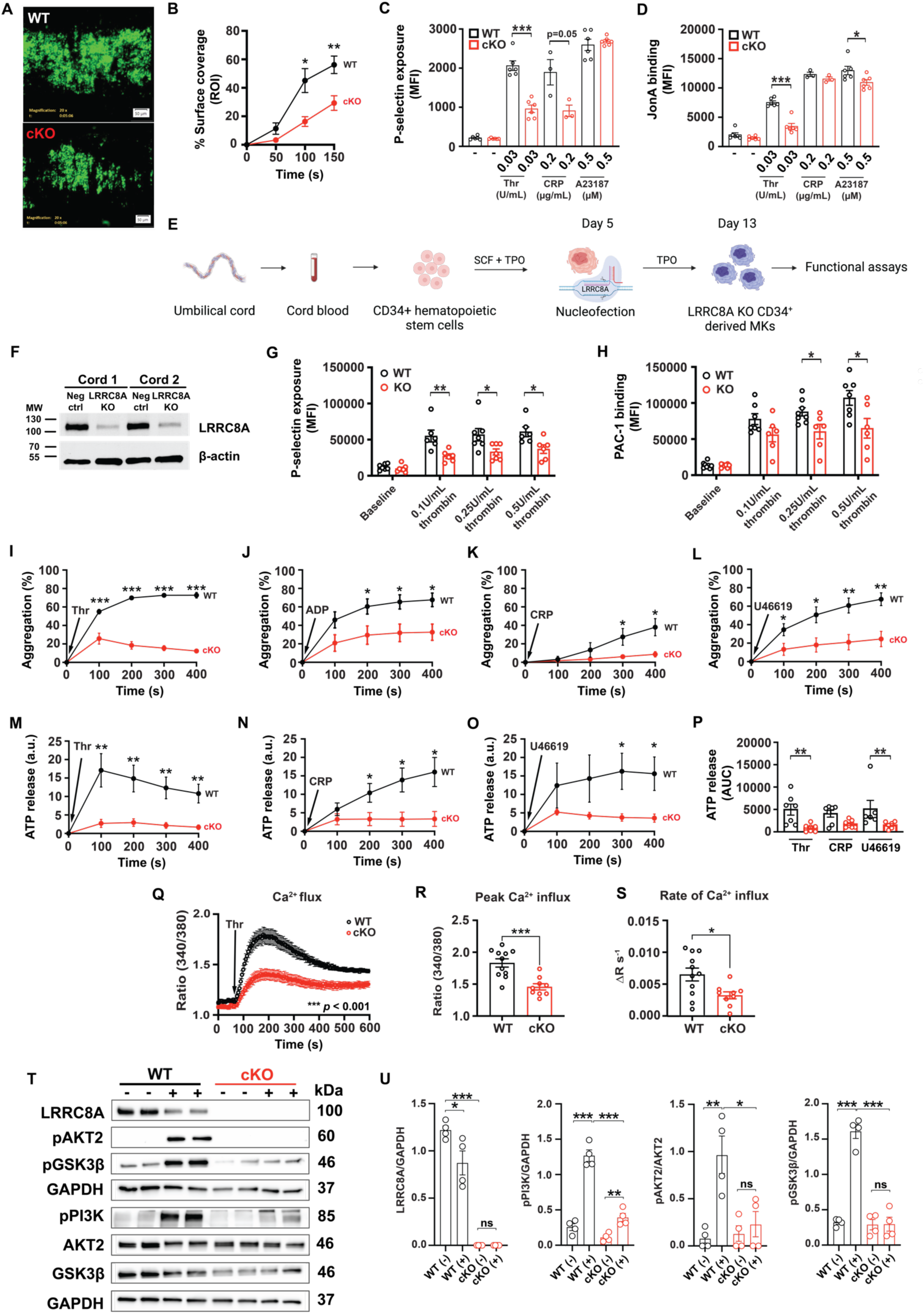
LRRC8A deletion impairs agonist-induced platelet adhesion, activation, aggregation, Ca^2+^ signaling and PI3K-AKT-GSK3β signaling. **a**, **b** Adhesion of platelets to a fibrillar collagen-coated surface under shear conditions (650 s^−1^) in whole blood isolated from *Lrrc8a^fl/fl^* (WT, n = 6, *left*) and *Pf4-Cre;Lrrc8a^fl/fl^*(cKO, n = 5, *right*) mice, as seen by fluorescence microscopy (Alexa Fluor^®^ 488). Quantification data from platelets from WT (n = 6) and cKO (n = 5) mice are shown in **b**. **c, d** P-selectin exposure (**c**) and JonA binding to activated αIIbβ3 integrin (**d**) as measured by flow cytometry of platelets isolated from WT and cKO mice and stimulated with thrombin (0.03 U/mL, n = 6 per group), CRP (0.2 µg/mL, n = 3), or the calcium ionophore A23187 (0.5 µM, n = 6 per group). **e** Illustration depicting the generation of LRRC8A KO CD34^+^-derived megakaryocytes from CD34^+^ hematopoietic stem cells isolated from umbilical cord blood. **f** LRRC8A expression in CRISPR-edited LRRC8A KO CD34^+^-derived human megakaryocytes generated from two independent cord blood samples. **g** Activation of CRISPR-edited LRRC8A KO CD34^+^-derived human megakaryocytes as determined by P-selectin exposure following thrombin stimulation at 0.1 U/mL, 0.25 U/mL or 0.5 U/mL for 15 min. **h** Activation of CRISPR-edited LRRC8A KO CD34^+^-derived human megakaryocytes as determined by PAC-1 binding to activated αIIbβ3 integrin following thrombin stimulation at 0.1 U/mL, 0.25 U/mL or 0.5 U/mL for 15 min. **i**, **j**, **k**, **l** Platelet aggregometry from WT and cKO mice stimulated with thrombin (**i**, 0.05 U/mL, n = 7 per group), ADP (**j**, 10 – 20 μM ADP and 30 µg/mL fibrinogen, n = 6 per group), CRP (**k**, 0.05 – 0.15 μg/mL, n = 7 per group), and U46619 (**l**, 0.5 – 0.8 μM, n = 6 per group). Concurrent ATP release from platelets treated with thrombin, CRP, and U46619 shown in **m**, **n** and **o**, respectively. Area under curve (AUC) of ATP release are shown in **p**. **q**, **r**, **s** Cytosolic Ca^2+^ measured using the ratiometric dye Fura-2 in platelets isolated from WT (n = 11) and cKO (n = 9) mice stimulated with thrombin (0.02 U/mL) for 5 min (**q**), with corresponding measurements of peak Ca^2+^ (**r**) and rate of Ca^2+^ rise (**s**). **t** Western blots detecting LRRC8A; pPI3K^Tyr458^; pAKT2^Ser474^; AKT2; pGSK-3β^Ser9^; GSK-3β; and GAPDH in platelets from WT and cKO mice following stimulation with PAR4-AP (400 µM) for 5 min during aggregometry, with densitometric quantification of LRRC8A / GAPDH; pPI3K / GAPDH; pAKT2 / AKT2 and pGSK-3β / GAPDH shown in **u**. Data are represented as mean ± S.E.M. Statistical significance was determined by unpaired T-test for **b** – **d**, **g** – **o**, **r**, **s**, and **u**, and by two-way ANOVA for **q**. For **p**, statistical significance was determined by unpaired T-test (Thr) or Mann-Whitney test (CRP, U46619). * *p* < 0.05; ** *p* < 0.01; *** *p* < 0.001. Figure **e** was made with BioRender.com

Since the functional defects observed in LRRC8A-null platelets in response to various agonists suggest impairments in common signaling pathways, we next measured cytosolic Ca^2+^ levels in response to thrombin (**Fig. 2Q-S**), and PI3K-AKT signaling following PAR4 stimulation (**Fig. 2T-U**), in WT and LRRC8A-null platelets. LRRC8A deletion lowers peak cytosolic Ca^2+^ levels (**Fig. 2Q&R**) and rate of rise of intracellular Ca^2+^ (**Fig. 2S**) in thrombin-stimulated platelets -indicating impaired Ca^2+^ signaling. This is also associated with marked reductions in PI3K-AKT2 signaling in LRRC8A-null platelets stimulated with PAR4-AP as assessed by pPI3K, pAKT2, pGSK3β, (**Fig. 2T&U**) and in the MEG-01 human megakaryoblast cell line after shRNA-mediated LRRC8A KD followed by thrombin stimulation (**Supplementary** Fig. 3A**&B**). Collectively, these data from both murine platelets and hiPSC MKs reveal that LRRC8A regulates common signaling pathways in platelet activation.

### LRRC8A deletion attenuates arteriolar and arterial thrombosis

To determine the pathophysiological function of platelet LRRC8A in thrombosis, we used WT and LRRC8A cKO mice in mouse models of laser-induced cremaster arteriolar thrombosis (**Fig. 3A-C**) and FeCl_3_-induced carotid arterial thrombosis (**Fig. 3D&E**). We also examined hemostasis by measuring bleeding times (**Fig. 3F**) and blood loss following amputation of the tail tip (**Fig. 3G**). Deletion of platelet-specific LRRC8A significantly reduces platelet thrombus formation compared to WT mice after laser-induced cremaster arteriolar injury (**Fig. 3A-C**, **Video S3&4**). Consistent with this finding, the time to occlusion (TTO) in a FeCl_3_-injured carotid artery is markedly prolonged in cKO mice compared to WT mice (**Fig. 3D&E**). Notably, 5/7 cKO mice had TTO greater than the 30 min (1,800 sec) time limit of the experiment. Despite reduced platelet thrombus formation in the cKO mice, no differences were observed in tail bleeding times and the amount of blood loss observed at the site of tail tip amputation (**Fig. 3F-G**). These results suggest that platelet LRRC8A contributes to the pathology of arterial thrombosis, but not hemostasis, in mice.

**Figure 3.**
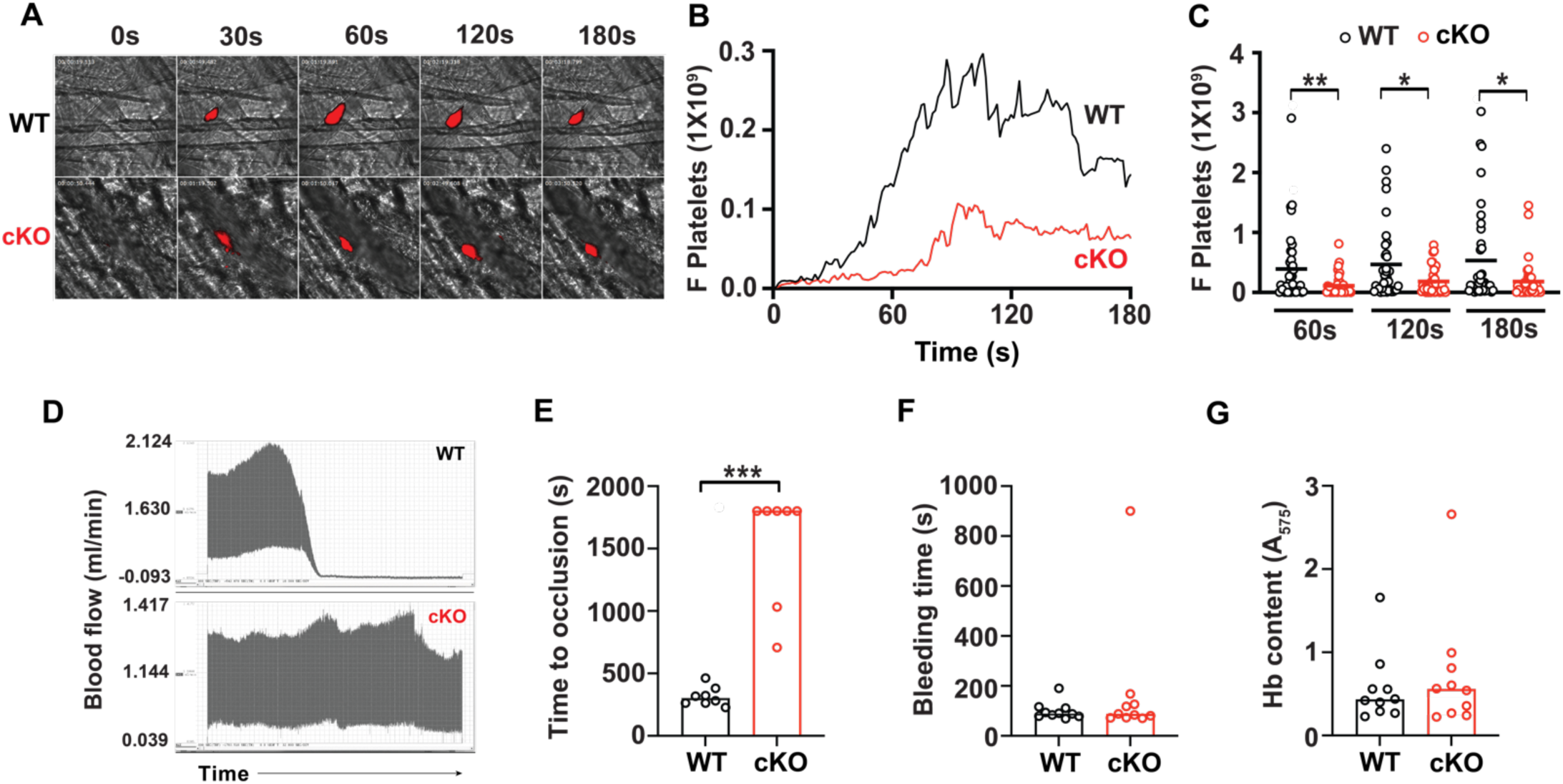
Platelet-targeted LRRC8A deletion impairs injury-induced arterial thrombosis, without prolonging hemostasis. **a, b,c** Arterial thrombosis in the cremasteric artery of *Lrrc8a^fl/fl^* (WT) and *Pf4-Cre;Lrrc8a ^fl/fl^* (cKO) mice following laser-induced injury as observed by real-time confocal intravital microscopy (*Red* = DyLight 649 anti-CD42C) (**a**) with median time course shown in **b** and quantification of the fluorescent reporter at 60, 120 and 180 seconds post-injury shown in **c** (n = 37 for both WT and cKO). **d** Representative traces of arterial blood flow through the carotid artery of WT (*top*) and cKO (*bottom*) mice following FeCl_3_-induced injury. **e** Time taken for the carotid artery of WT (n = 8) and cKO (n = 7) mice to occlude following FeCl_3_-induced injury**. f** Total bleeding time following tail tip amputation of WT (n = 10) and cKO (n = 10) mice, with corresponding hemoglobin concentrations of red blood cell lysates collected during the assay shown in **g**. Data in **c** are represented as mean only, whereas data in **b**, **e**, **f** and **g** are represented as median. Statistical significance was determined by Mann-Whitney test for **c** and **e**, and by unpaired T-test for **f** and **g**. * *p* < 0.05; ** *p* < 0.01; *** *p* < 0.001.

### The LRRC8 complex amplifies platelet activation and aggregation by contributing stimulatory ATP and ADP

As the LRRC8 channel complex forms an anion channel, we asked how loss of an anion conductance might impair Ca^2+^ signaling (**Fig. 2Q-S**). Based on our observations that ATP release is abrogated in LRRC8A KO platelets in response to multiple agonists (**Fig. 2M-P**), and the finding that LRRC8A can form an ATP release channel ^41,54^, we hypothesized that platelet LRRC8 channels contribute to local ATP efflux to amplify platelet activation and aggregation. We first directly measured swell-activated ATP currents in WT MKs by comparing the hypotonicity-induced inward current in cells where all intracellular anions were replaced with either 1 mM ATP^4-^ (**Fig. 4A, D&G**) or 50 mM ATP^4-^ (**Fig. 4B, E&G**). MKs dialyzed with 50 mM ATP^4-^ as the only permeant anion yield robust SN-401 (DCPIB)-sensitive inward ATP currents, indicative of ATP efflux, 4.4-fold larger than cells dialyzed with 1 mM ATP^4-^, and these ATP currents are abolished in LRRC8A KO MKs (**Fig. 4C, F&G**), demonstrating LRRC8A-dependent ATP currents in response to MK volume changes. Similar results were obtained using Ad-shSCR and Ad-shLRRC8A transduced MEG-01 cells (**Supplementary Fig. 4**).

**Figure 4.**
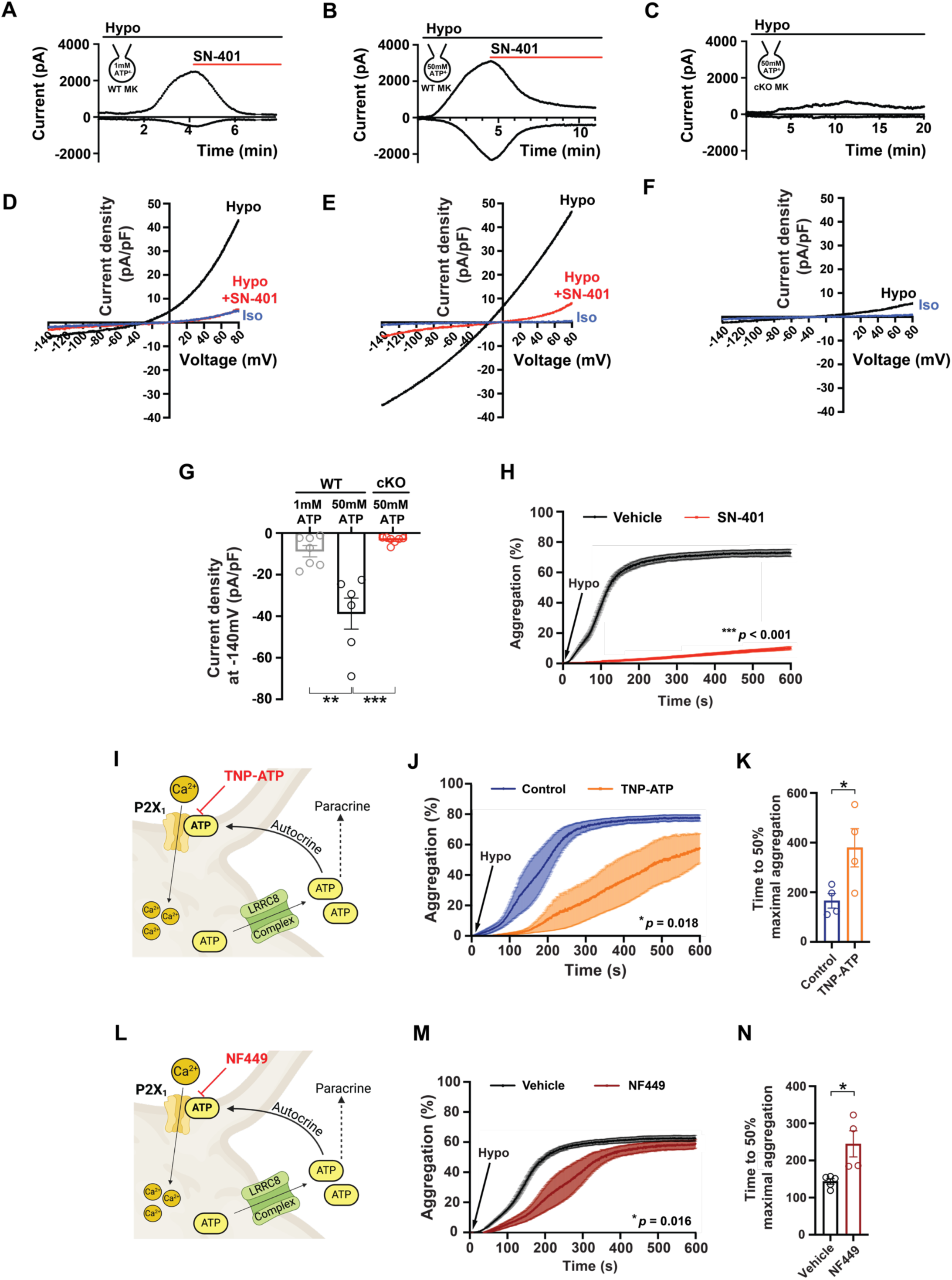
Platelet LRRC8 channels permeate ATP and potentiate platelet aggregation via autocrine and paracrine P2X1 receptor activation. **a – c** Current-time relationship of inward I_ATP_ (ATP efflux) and outward VRAC in MKs isolated from *Lrrc8a^fl/fl^* (WT) mice induced by hypotonic (210 mOsm) swelling with an intracellular ATP concentration of 1 mM (**a**) or 50 mM (**b**) followed by application of 10 µM SN-401 (DCPIB), and in MKs isolated from *Pf4-Cre;Lrrc8a ^fl/fl^* (cKO) mice (**c**) with an intracellular ATP concentration of 50 mM. The inward component of the current represents I_ATP_ generated from ATP efflux. Respective current-voltage relationships of inward I_ATP_ (ATP efflux) and outward VRAC elicited from voltage ramps from -140 mV to +80 mV are shown in **d** – **f**. **g** Mean current densities of inward I_ATP_ (ATP efflux) at -140 mV in MKs isolated from WT mice after hypotonic swelling, with an intracellular concentration of 1 mM ATP (n = 7) or 50 mM ATP (n = 6), and in MKs isolated from cKO mice with an intracellular concentration of 50 mM ATP (n = 7). **h** Aggregometry of human platelets upon switching from isotonic (270 mOsm) to hypotonic (180 mOsm) solution in the presence of vehicle (0.02% DMSO; n = 6, isolated from one healthy volunteer) or 10 µM SN-401 (DCPIB; n = 5, isolated from one healthy volunteer). **i** Working model of platelet LRRC8 channel ATP efflux and subsequent autocrine and paracrine activation of the ATP-gated P2X1 Ca^2+^ permeant channel, and P2X1 channel inhibition by TNP-ATP. **j, k** Aggregometry of human platelets activated by decreasing HTB osmolarity in the presence (n = 4, isolated from one healthy volunteer) or absence (n = 4, isolated from one healthy volunteer) of TNP-ATP (850 nM) – an inhibitor of P2X1 (**i**). Time to reach 50% maximal aggregation are shown in **k**. **l** Working model of platelet LRRC8 channel ATP efflux and subsequent autocrine and paracrine activation of the ATP-gated P2X1 Ca^2+^ permeant channel, and P2X1 channel inhibition by NF449. **m, n** Aggregometry of human platelets activated by decreasing HTB osmolarity in the presence (n = 4, isolated from one healthy volunteer) or absence (n = 5, isolated from one healthy volunteer) of NF449 (500 nM) – an inhibitor of P2X1 (**l**). Time to reach 50% maximal aggregation are shown in **n**. Data are represented as mean ± S.E.M. Statistical significance was determined by unpaired T-test for **g**, **k** and **n**, and by two-way ANOVA for **h**, **j** and **m**. * *p* < 0.05; ** *p* < 0.01; *** *p* < 0.001.

Given that hypotonic swelling activates LRRC8 channels in MKs, the precursors to platelets, we next asked if platelet aggregation can be induced by hypotonic swelling in the absence of a traditional agonist. Human platelets aggregate robustly when osmolarity is reduced from 270 mOsm to 180 mOsm (hypotonic stimulation), and this is robustly inhibited by SN-401 (**Fig. 4H**). As ATP amplifies platelet activation and aggregation by activating the calcium-permeable ATP-gated P2X1 receptor ^16^, we next examined the effects of two P2X1 receptor antagonists, TNP-ATP (**Fig. 4I**) and NF449 (**Fig. 4L**)^17^, on platelet aggregation induced by hypotonic swelling. Both TNP-ATP (**Fig. 4J**) and NF449 (**Fig. 4M**) prolong the time to 50% maximal aggregation in response to hypotonic swelling, but do not impair maximal aggregation (**Fig. 4J&K**; **Fig 4M&N**). These results demonstrate that swelling-induced platelet LRRC8 ATP release accelerates the kinetics of platelet aggregation via autocrine and paracrine P2X1 activation.

However, this mechanism does not account for the level of suppression of platelet aggregation by LRRC8 inhibition upon hypotonic swelling (**Fig. 4H**), suggesting that reduced LRRC8 mediated ATP efflux and diminished P2X1 receptor activation cannot alone account for impairments in platelet function in LRRC8A KO mice. Since ADP is also a platelet activating anion released from platelets to induce platelet aggregation ^55^, we asked whether LRRC8 complexes can permeate ADP, in addition to ATP. Accordingly, we first directly measured swell-activated ADP currents in WT MKs by comparing the hypotonic-induced inward current in cells where all intracellular anions were replaced with either 1 mM ADP^3-^ (**Fig. 5A, D&G**) or 50 mM ADP^3-^ (**Fig. 5B, E&G**). When MKs are dialyzed with 50 mM ADP^3-^ as the only permeant anion, MKs yield robust SN-401 (DCPIB)-sensitive inward currents, indicative of ADP efflux, 2.1-fold larger than cells dialyzed with 1 mM ADP^3-^, and these ADP currents are abolished in LRRC8A KO MKs (**Fig. 5C, F&G**), demonstrating LRRC8A-dependent ADP currents in response to MK volume changes. Similar results were obtained using Ad-shSCR and Ad-shLRRC8A transduced human MEG-01 cells (**Supplementary Fig. 5**). ADP activates platelets via two G-protein coupled receptors, P2Y_1_ and P2Y_12_ (**Fig. 5H**) ^56^. We first examined the contribution of P2Y_12_ stimulation by ADP released from swell-activated LRRC8 channels by applying ticagrelor, a clinically used P2Y_12_ receptor inhibitor. Remarkably, ticagrelor fully suppressed swell-induced LRRC8 mediated platelet aggregation (**Fig. 5I**). Since ADP-mediated P2Y_12_ receptor stimulation subsequently activates the PI3K-AKT axis to promote dense granule secretion and entry into the irreversible phase of aggregation ^6,57,58^, we examined AKT signaling in response to hypotonic swelling +/-P2Y_12_ receptor inhibition with ticagrelor. Swelling-induced phosphorylation of platelet AKT1, and GSK-3β (a known target of AKTs and a regulator of platelet function^59^), are significantly reduced by ticagrelor (**Supplementary Fig. 6**) . To evaluate the contributions of P2Y_1_ to LRRC8-mediated platelet aggregation we applied MRS2179 ^60,61^, a selective P2Y_1_ inhibitor, to hypotonically-stimulated platelets (**Fig. 5J**). Hypotonic swelling-induced platelet aggregation in the presence of the P2Y_1_ inhibitor MRS2179 ^60,61^ delays the time to 50% maximal aggregation (**Fig. 5K**), but not maximal aggregation, like results obtained for P2X1 inhibition by TNP-ATP and NF449 (**Fig. 4I-N**). Additionally, whilst NF449 is a selective antagonist of the P2X1 receptor at lower concentrations, concurrent inhibition of both P2X1 and P2Y_1_ receptors occurs when used at higher concentrations ^62^. Dual inhibition of P2X1 and P2Y_1_ using high concentration (50 µM) NF449, also suppresses hypotonic swelling-induced platelet aggregation (**Fig. 5L&M**). Whilst P2X1 mediates Ca^2+^ influx across the plasma membrane, P2Y_1_ activation results in the mobilization of intracellular Ca^2+^ stores ^56^, suggesting that combined P2X1/P2Y_1_ inhibition using high concentration NF449 diminishes both Ca^2+^ signaling mechanisms triggered by LRRC8-mediated aggregation. Taken together, these data are consistent with a model in which agonist-induced shape change activates platelet LRRC8 channels to efflux ATP and ADP, thereby amplifying the central PI3K-AKT and Ca^2+^ signaling pathways via activation of P2X1, P2Y_1_, and P2Y_12_ receptors.

**Figure 5.**
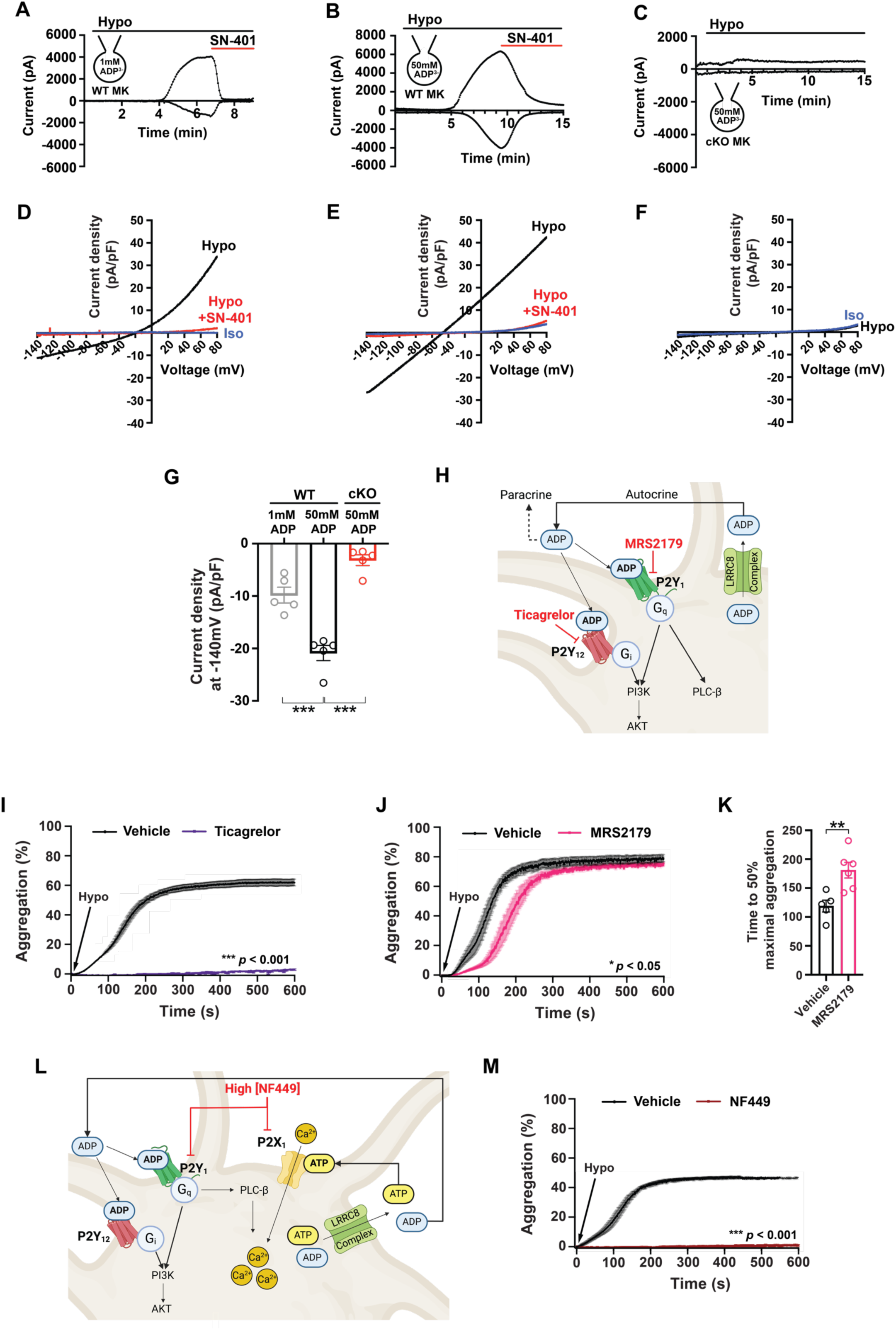
Platelet LRRC8 channels permeate ADP and potentiate platelet aggregation via autocrine/paracrine P2Y_1_ and P2Y_12_ receptor activation. **a – c** Current-time relationship of inward I_ADP_ (ADP efflux) and outward VRAC in MKs isolated from *Lrrc8a^fl/fl^* (WT) mice induced by hypotonic (210 mOsm) swelling with an intracellular ADP concentration of 1 mM (**a**) or 50 mM (**b**) followed by application of 10 µM SN-401 (DCPIB), and in MKs isolated from *Pf4-Cre;Lrrc8a ^fl/fl^*(cKO) mice (**c**) with an intracellular ADP concentration of 50 mM. The inward component of the current represents I_ADP_ generated from ADP efflux. Respective current-voltage relationships of inward I_ADP_ (ADP efflux) and outward VRAC elicited from voltage ramps from -140 mV to +80 mV are shown in **d** – **f**. **g** Mean current densities of inward I_ADP_ (ADP efflux) at -140 mV in MKs isolated from WT mice after hypotonic swelling with an intracellular concentration of 1 mM ADP (n = 5), 50 mM ADP (n = 5), and in MKs isolated from cKO mice after hypotonic swelling with an intracellular concentration of 50 mM ADP (n = 5). **h** Working model of platelet LRRC8 channel ADP efflux, subsequent autocrine and paracrine activation of P2Y_1_ and P2Y_12_ receptors, and inhibition by ticagrelor (P2Y_12_) and MRS2179 (P2Y_1_). **i** Aggregometry of human platelets following the addition of molecular grade water to decrease HTB osmolarity from isotonic (270 mOsm) to hypotonic (180 mOsm) in the presence (n = 5, isolated from one healthy volunteer) or absence (n = 5, isolated from one healthy volunteer) of ticagrelor (85 nM) – a P2Y_12_ inhibitor. **j** Aggregometry of human platelets activated by decreasing HTB osmolarity in the presence (n = 6, isolated from one healthy volunteer) or absence (n = 5, isolated from one healthy volunteer) of MRS2179 (700 nM) – a P2Y_1_ inhibitor. Time to reach 50% maximal aggregation are shown in **k**. **l** Working model of platelet LRRC8 channel ATP and ADP efflux, subsequent autocrine and paracrine activation of P2X1 (ATP) and P2Y_1_ / P2Y_12_ (ADP) receptors. NF449 at 50 µM inhibits both P2X1 and P2Y_1_ receptors. **m** Aggregometry of human platelets activated by decreasing HTB osmolarity in the presence (n = 5, isolated from one healthy volunteer) or absence (n = 4, isolated from one healthy volunteer) of 50 µM NF449 inhibiting both P2X1 and P2Y_1_ receptors. Data are represented as mean ± S.E.M. Statistical significance was determined by unpaired T-test for **g** and **k**, and by two-way ANOVA for **i**, **j**, and **m**. * *p* < 0.05; ** *p* < 0.01; *** *p* < 0.001.

### Small-molecule LRRC8 complex inhibitors block MK ATP/ADP efflux, and platelet activation, signaling and aggregation

To investigate the therapeutic potential of targeting the LRRC8 channel, we examined whether established small-molecule inhibitors, SN-406 ^51^ (**Fig. 6A**), and SN-407 ^51^ (**Supplementary Fig. 7A**) inhibit platelet function. Compared to SN-401, SN-406 and SN-407 are more potent VRAC inhibitors with IC_50_ measured in HEK cells of 3.1 and 1.6 µM respectively, as compared to 3.9 µM for SN-401 ^51^. Indeed, SN-406 robustly inhibits swell-activated ATP (**Fig. 6B-D**) and ADP (**Fig. 6E-G**) efflux in MEG-01 cells. Comparable results are obtained when using SN-407 (**Supplementary Fig. 7B-G**).

**Figure 6.**
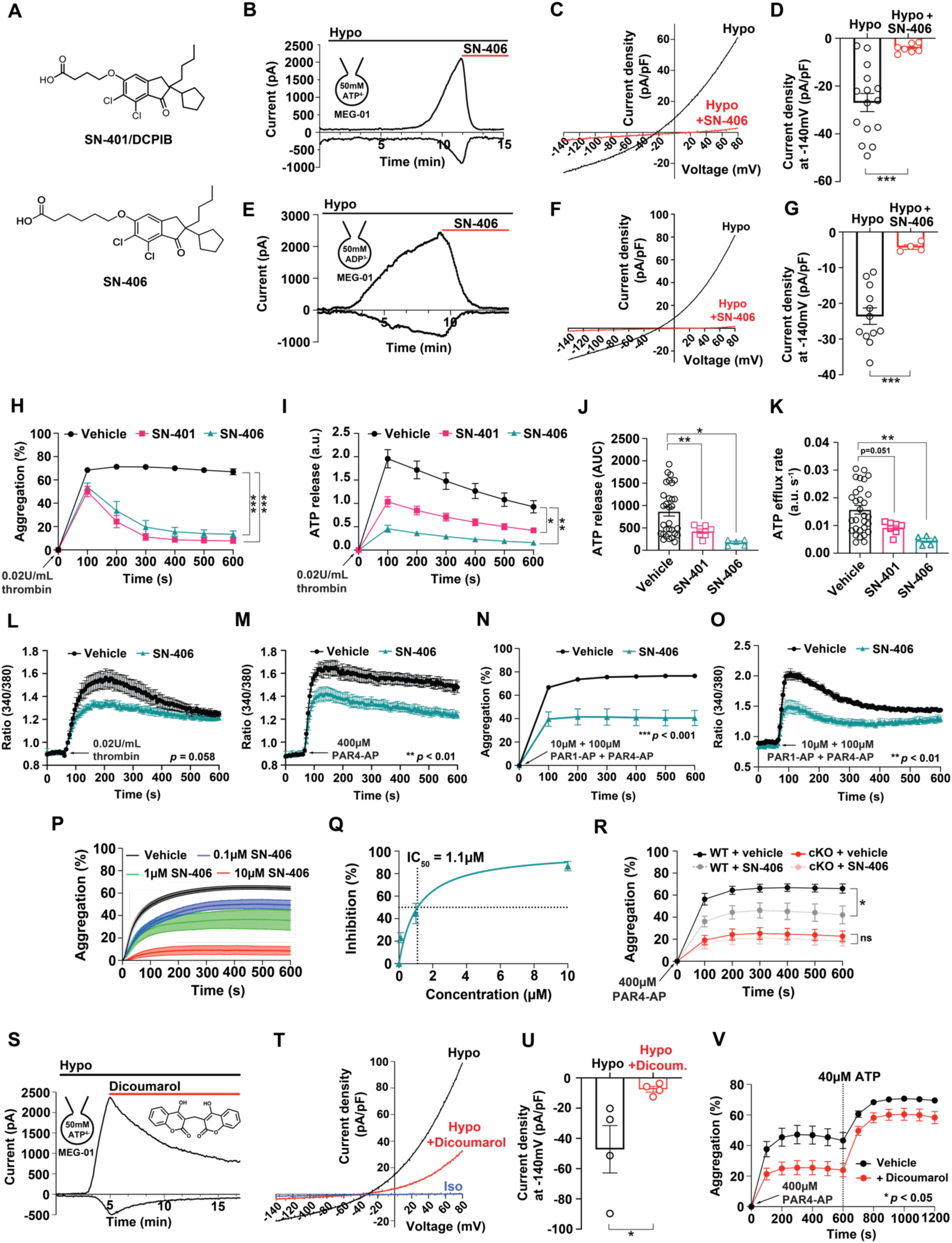
LRRC8 channel small molecule inhibitors suppress agonist-induced aggregation, ATP release, and calcium influx. **a** Molecular structure of the small molecule LRRC8 channel complex modulator SN-401 (DCPIB) and SN-406 – a derivative of SN-401. **b** Current-time relationship of inward I_ATP_ (ATP efflux) and outward VRAC induced by hypotonic swelling in MEG-01 cells and subsequent inhibition by application of 10 µM SN-406. **c, d** Current-voltage relationship of inward I_ATP_ (ATP efflux) and outward VRAC during voltage ramps from -140 mV to +80 mV in MEG-01 cells after hypotonic swelling in the absence or presence of 10 µM SN-406, with mean current densities of inward I_ATP_ at -140 mV shown in **d** (n = 7 – 15). **e** Current-time relationship of inward I_ADP_ (ADP efflux) and outward VRAC induced by hypotonic swelling in MEG-01 cells and subsequent inhibition by application of 10 µM SN-406. **f, g** Current-voltage relationship of inward I_ADP_ (ADP efflux) and outward VRAC during voltage ramps from -140 mV to +80 mV in MEG-01 cells after hypotonic swelling in the absence or presence of 10 µM SN-406, with mean current densities of inward I_ADP_ at -140 mV shown in **g** (n = 4 – 12). **h – k** Aggregometry of platelets isolated from WT C57BL/6J mice stimulated with thrombin (0.02 U/mL) in the presence of vehicle (0.02% DMSO; n = 37), SN-401 (10 µM; n = 7), or SN-406 (10 µM, n = 5), with concurrent ATP release (n = 5 – 30) shown in **i**, **j** and **k**. **l, m** Cytosolic calcium measured using the ratiometric dye Fura-2 in platelets isolated from WT C57BL/6J mice stimulated with thrombin (**l;** 0.02 U/mL; n = 8) or PAR4-AP (**m**; 400 µM; n = 8) in the presence of vehicle only (0.02% DMSO) or SN-406 (10 µM). **n, o** Aggregometry of human platelets stimulated with PAR1-AP and PAR4-AP (10 µM and 100 µM, respectively) in the presence of vehicle (0.02% DMSO; n = 10 from three healthy volunteers) or SN-406 (10 µM; n = 11 from three healthy volunteers), with measurements of cytosolic calcium shown in **o**. **p** Aggregometry of PAR4-AP (400 µM) stimulated mouse platelets in the presence of vehicle (0.02% DMSO; n = 35), and SN-406 at 0.1 µM (n = 5), 1 µM (n = 5), or 10 µM (n = 6). **q** SN-406 dose-dependent inhibition of PAR4-AP (400 µM) stimulated mouse platelet aggregation (SN-406: 0 – 10 µM; n = 5 – 6). **r** Aggregometry of platelets isolated from *Pf4-Cre;Lrrc8a ^fl/fl^* (cKO) and *Lrrc8a^fl/fl^* (WT) littermate controls stimulated by PAR4-AP (400 µM) in the presence of vehicle (0.02% DMSO; n = 5 for WT, n = 7 for cKO) or 1.1 µM SN-406 (n = 6 for WT, n = 8 for cKO). **s** Current-time relationship of inward I_ATP_ and outward VRAC induced by hypotonic (210 mOsm) swelling in MEG-01 cells and subsequent inhibition by application of 10 µM dicoumarol. **t**, **u** Current-voltage relationship of inward I_ATP_ and outward VRAC during voltage ramps from -140 mV to +80 mV in MEG-01 cells after hypotonic swelling in the absence or presence of 10 µM dicoumarol, with mean current densities of inward I_ATP_ -140 mV shown in **u** (n = 4 for both groups). **v** Aggregometry of platelets isolated from WT C57BL/6J mice stimulated with PAR4-AP (400 µM) in the presence of vehicle (0.1% DMSO; n = 4) or dicoumarol (20 µM; n = 3), with subsequent application of ATP (40 µM). Data are represented as mean ± S.E.M. Statistical significance was determined by unpaired T-test for **d**, **g**, **j**, **k** and **u**, and by two-way ANOVA for **h**, **i**, **l**, **m**, **n**, **o**, **r** and **v**. * *p* < 0.05; ** *p* < 0.01; *** *p* < 0.001.

Consistent with these effects on LRRC8-mediated ATP currents in patch-clamp studies, SN-401 and SN-406 inhibit thrombin-stimulated platelet aggregation (**Fig. 6H**) and markedly reduce total ATP release (**Fig. 6I&J**) and ATP efflux rate (**Fig. 6K**), as compared to vehicle. In addition, SN-406 and SN-407 significantly inhibit platelet aggregation, ATP release, and ATP efflux rate after PAR4-stimulation (**Supplementary Fig. 7H-K**). Also, as observed in LRRC8A KO platelets, SN-406 treatment reduces both thrombin-(**Fig. 6L**) and PAR4-AP-(**Fig. 6M**) stimulated Ca^2+^ flux, consistent with a mechanism of impaired ATP-stimulated Ca^2+^ signaling. In addition, SN-406 significantly suppresses human platelet aggregation in response to PAR1+PAR4 stimulation (**Fig. 6N**) and impairs agonist-induced Ca^2+^ signaling (**Fig. 6O**), as observed in mouse platelets. SN-406 exhibits a clear dose-response relationship in PAR4-AP-induced mouse platelet aggregation (**Fig. 6P**), yielding an IC_50_ of 1.1 µM (**Fig. 6Q**), which is comparable to the IC_50_ of 3.1 µM for VRAC inhibition in HEK cells published previously ^51^. Applying 1.1 µM SN-406 to WT and LRRC8A KO platelets shows significant suppression of platelet aggregation in WT platelets with no effect on LRRC8A KO platelets, consistent with SN-406-LRRC8 complex on-target activity (**Fig. 6R**). We next tested the VRAC inhibitor dicoumarol (**Fig. 6S**), which is structurally distinct from the SN-40X compounds optimized off the SN-401 (DCPIB) backbone. Dicoumarol, identified in a small molecule screen, was recently shown to inhibit LRRC8A-mediated ATP release from microglia ^41^. We find dicoumarol also inhibits swell-induced ATP efflux in MEG-01 cells (**Fig. 6S-U**). When applied to murine platelets, similar to SN-40X compounds, dicoumarol markedly suppresses platelet aggregation, and this is rescued by exogenous ATP (**Fig. 6V**).

Through further compound optimization, we developed SN-89B, a more potent LRRC8 inhibitor, with lower predicted plasma membrane protein binding than those previously described. SN-89B strongly inhibits ATP and ADP efflux elicited in response to hypotonic swelling in MEG-01 cells (**Fig. 7A-D** and **Supplementary Fig. 8**). In human platelets, SN-89B inhibits aggregation in a dose-dependent manner (**Fig. 7E**), with an IC_50_ of 840 nM (**Fig. 7F**). In agonist-stimulated human and mouse platelets, platelet activation is fully suppressed by SN-89B as assessed by P-selectin exposure (**Fig. 7G&I**) and αIIbβ3 activation (**Fig. 7H&J**). Furthermore, SN-89B significantly reduces agonist-induced human platelet aggregation (**Fig. 7K**), as well as phosphorylation of AKT1, AKT2, and GSK-3β (**Fig. 7L&M**), which are components of the PI3K-AKT axis. Following these *in vitro* studies, we then proceeded to treat WT mice with vehicle or 50 mg/kg SN-89B intraperitoneally for 8 days to determine its effects on thrombosis *in vivo* using the FeCl_3_-induced carotid arterial thrombosis model (**Fig. 7N**). SN-89B significantly prolongs FeCl_3_-induced carotid arterial thrombosis (**Fig. 7O**) as compared to the vehicle control. Taken together, these data demonstrate LRRC8 channels as ATP/ADP release channels that regulate P2X1, P2Y_1_ and P2Y_12_ mediated Ca^2+^ and PI3K-AKT signaling to amplify platelet activation, adhesion and aggregation and *in vivo* thrombosis, in addition to demonstrating the pharmacological tractability of targeting LRRC8 channels as a novel anti-thrombotic strategy.

**Figure 7.**
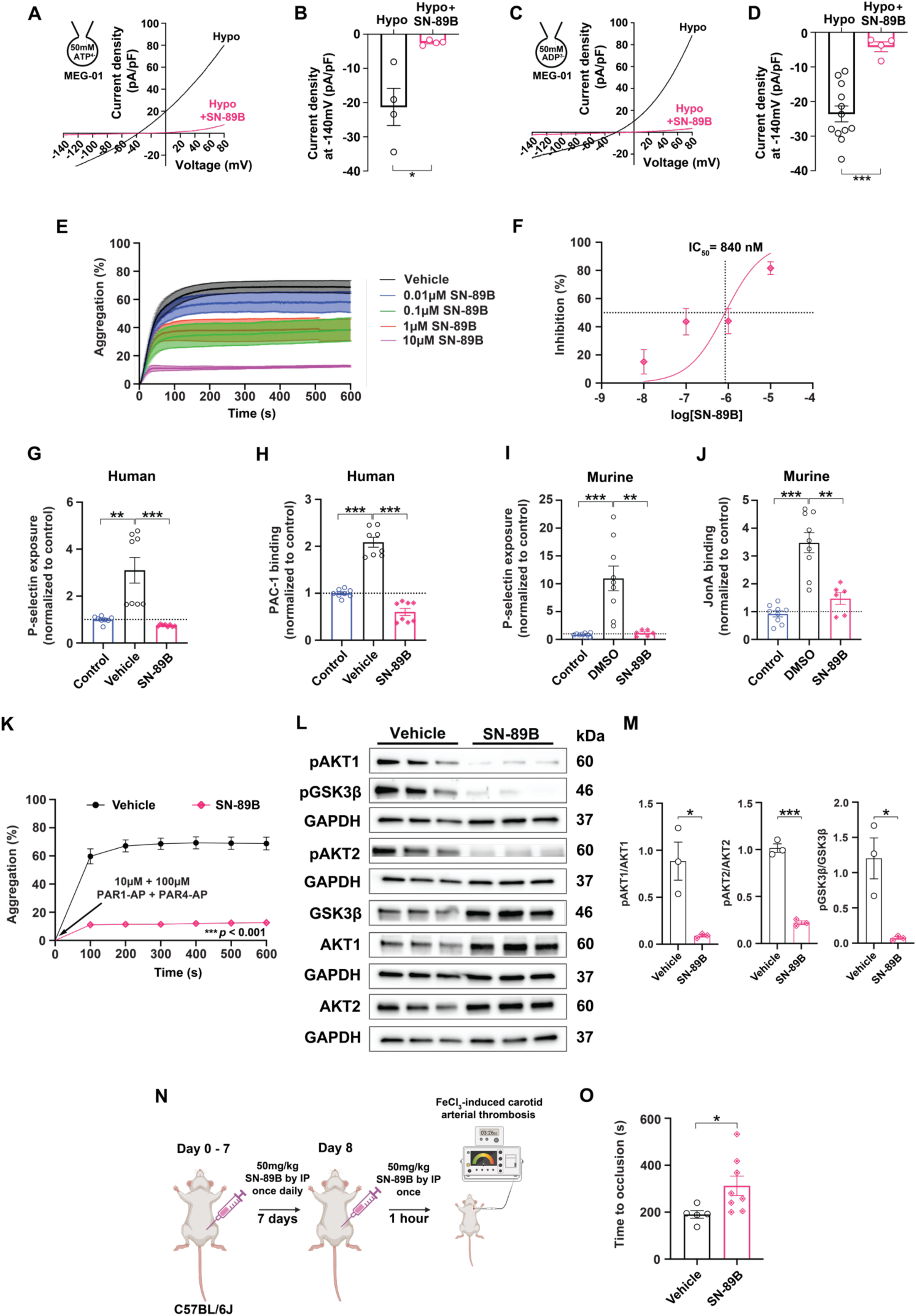
LRRC8 channel inhibitor SN-89B suppresses agonist-induced platelet activation, aggregation, PI3K-AKT-GSK3β signaling and *in vivo* thrombosis. **a –d** Current-voltage relationship of inward I_ATP_ (ATP efflux; a), I_ADP_ (ADP efflux; b), and outward VRAC (both **a** and **b**) during voltage ramps from -140 mV to +80 mV in MEG-01 cells after hypotonic swelling in the absence or presence of 10 µM SN-89B. Mean current densities of inward I_ATP_ and I_ADP_ at -140 mV are shown in **b** (n = 4) and **d** (n = 4 – 12), respectively. e, f Aggregometry of PAR1-AP and PAR4-AP (10 µM and 100 µM, respectively) stimulated human platelets in the presence of vehicle (0.02% DMSO; n = 6) and SN-89B at either 0.01 µM (n = 3), 0.1 µM (n = 3), 1 µM (n = 4), or 10 µM (n = 4). SN-89B dose-dependent inhibition of PAR1-AP and PAR4-AP-stimulated human platelet aggregation (SN-89B: 0 -10 µM; n = 3 – 6, isolated from one healthy volunteer) is shown in **f. g, h** P-selectin exposure (**g**) and PAC-1 binding to activated αIIbβ3 integrin (h) as measured by flow cytometry of human platelets stimulated with PAR1-AP and PAR4-AP (10 µM and 100 µM, respectively) in the presence of vehicle (0.02% DMSO) or SN-89B (10 µM). Each sample was normalized to the average of unstimulated controls (n = 8 for all groups, isolated from two healthy volunteers). **i, j** P-selectin exposure (i) and JonA binding to activated αIIbβ3 integrin (j) as measured by flow cytometry of mouse platelets stimulated with PAR4-AP (400 µM) in the presence of vehicle (0.02% DMSO, n = 9) or SN-89B (10 µM, n = 6). Each sample was normalized to the average of unstimulated controls (n = 9). **k** Aggregometry of human platelets stimulated with PAR1-AP and PAR4-AP (10 µM and 100 µM, respectively) in the presence of vehicle (0.02% DMSO, n = 6, isolated from one healthy volunteer) or SN-89B (10 µM, n = 4, isolated from one healthy volunteer). l Western blots detecting pAKT1^Ser473^; AKT1; pAKT2^Ser474^; AKT2; pGSK-3β^Ser9^; GSK-3β; and GAPDH in human platelets stimulated with PAR1-AP and PAR4-AP (10 µM and 100 µM, respectively), in the presence of vehicle (0.02% DMSO, n = 3, isolated from one healthy volunteer) or SN-89B (10 µM, n = 3, isolated from one healthy volunteer), with densitometric quantification of pAKT1 / AKT1; pAKT2 / AKT2 and pGSK-3β / GSK-3β shown in **m. n** Cartoon depicting the SN-89B dosing regimen used prior to FeCl_3_-induced carotid arterial thrombosis. o Time to occlusion of the carotid artery of WT C57BL/6J mice treated 50 mg/kg SN-89B (n = 8) or vehicle (n = 5) following FeCl_3_-induced injury. Data are represented as mean ± S.E.M. Statistical significance was determined by unpaired T-test for **b, d, g, h, i, j, m** and **o**, and by two-way ANOVA for **k.** * *p* < 0.05; ** *p* < 0.01; *** *p* < 0.001. Figure **n** was made with BioRender.com

## Discussion

Thrombotic diseases, including coronary artery disease, stroke, and pulmonary embolism, are the leading cause of death in the U.S. Platelets are the first blood cells recruited to the site of vascular injury. Adherent platelets undergo shape and volume changes that are early mechanical events in platelet activation and subsequently facilitate platelet aggregation^5^. However, the molecular mechanisms responsible for sensing and transducing these changes in platelet volume are unknown. As the initial findings of our study indicated LRRC8 proteins are highly expressed in human and mouse platelets and changes in LRRC8 proteins are associated with altered function (MPV and altered responses to several agonists), we hypothesized that LRRC8 channel complexes function as mechano-sensors linking platelet shape change to platelet activation. Collectively, our data from selective LRRC8A deletion in platelets, and human iPSC-derived MKs, reveal significant impairments in cell activation, adhesion, aggregation, and ATP release in response to multiple agonists, suggesting an LRRC8A-dependent defect in common intra-platelet signaling pathways. Consistent with this prediction, we observe impaired thrombin/PAR4-PI3K-AKT signaling, and reduced agonist-stimulated Ca^2+^ signaling -both common signaling pathways downstream of platelet agonist GPCRs, namely the ADP receptors P2Y_1_ and P2Y_12_, thrombin receptors PAR1 and PAR4, serotonin receptor 5-HT_2A_, and thromboxane A2 receptor TP^63^.

These observations led us to ask how activation of the LRRC8 *anion* channel might stimulate increases in both cytosolic Ca^2+^ and PI3K-AKT signaling? As the LRRC8 complex is an anion channel that also permeates larger anions such as ATP^54,64,65^ and has recently been shown to form an ATP release channel in microglia^41^, we hypothesized that platelet shape-change activates LRRC8-mediated ATP efflux, which subsequently activates ATP-gated P2X1 receptor^16,66^ Ca^2+^ influx in a paracrine/autocrine manner to amplify platelet activation and aggregation. In line with this mechanism, Tomasiak *et al*. showed that thrombin-induced ATP secretion from porcine platelets increases under hypoosmotic stimulation – conditions expected to activate LRRC8 channels^67^, and we find LRRC8A-dependent swell-activated ATP efflux can be directly measured in MKs in patch-clamp studies. Consistent with our model, pharmacological inhibition of the P2X1 receptor in our studies delayed rates of swelling-stimulated LRRC8-mediated aggregation. However, the fact that P2X1 inhibition did not diminish maximal aggregation as observed with pharmacological LRRC8 channel blockade, implicated other LRRC8 mediated molecular mechanisms.

Accordingly, we hypothesized that LRRC8 channels also permeate ADP, which stimulate P2Y_1_ and P2Y_12_ receptor signaling^56^ – the latter being a well-established therapeutic target of a number of clinically used anti-platelet agents, including clopidogrel, ticagrelor and prasugrel ^68^. Indeed, LRRC8A-dependent swell-activated ADP currents can be directly measured in MKs in patch-clamp studies. In platelets, P2Y_1_ inhibition delays swell-activated platelet aggregation, while P2Y_12_ inhibition robustly inhibits swell-activated platelet PI3K-AKT signaling and aggregation. We propose that LRRC8 complexes are activated by shape change during the early phases of agonist-induced aggregation which results in ATP/ADP release to amplify PI3K-AKT signaling via ADP-stimulated P2Y_12_ receptors, and Ca^2+^ signaling via ADP stimulated P2Y_1_ and ATP stimulated P2X1 receptors.

Interestingly, P2X1 receptor KO mice phenocopy platelet-targeted LRRC8A cKO mice with respect to platelet Ca^2+^ influx, adhesion, aggregation and thrombus formation^69^, with relatively preserved hemostasis in both P2X1 mice (30% increase in mean bleeding time)^69^ and LRRC8A cKO mice (0% increase in median, 70% increase in mean bleeding times). P2Y_1_ KO mice also exhibit impaired ADP-induced aggregation and impaired thrombus formation, but with 70 – 240% increases in mean bleeding times compared to respective controls ^70,71^, while P2Y_12_ KO mice exhibit *severely* prolonged mean bleeding times, 700 – 800% greater than controls ^72,73^. These comparatively smaller increases in bleeding time in P2X1 and P2Y_1_ KO mice are potentially due to decreased cytosolic calcium signaling but intact PI3K-AKT signaling, whereas ablation of P2Y_12_ fully abrogates PI3K-AKT signaling required for stable aggregation. Thus, pharmacological inhibition of LRRC8-mediated ATP/ADP efflux may provide a therapeutic strategy to tune P2X1, P2Y_1_ and P2Y_12_ receptor activation to dampen platelet aggregation and thrombosis, while minimally prolonging bleeding times, and maintaining hemostasis.

In line with this concept, VRAC inhibitors such as SN-401 (DCPIB) and more potent derivatives, SN-406 and SN-407^51^, strongly inhibit agonist-stimulated platelet aggregation, ATP release, and agonist-stimulated Ca^2+^ flux in both murine and human platelets Furthermore, SN-89B, a more potent SN-401 derivative with reduced predicted protein binding, strongly inhibited human platelet activation, agonist and hypotonic-induced aggregation, and PI3K-AKT signaling *in vitro*, and exerted antithrombotic properties *in vivo* – providing a proof-of-concept for developing small-molecule LRRC8 inhibitors as novel anti-platelet agents. Importantly, as SN-401, SN-406 and SN-407 have been previously demonstrated to improve glycemic control and insulin sensitivity in type 2 diabetes mouse models^51^, this class of molecules possess both anti-platelet and anti-diabetic properties which can be a desirable combination when treating patients with both diabetes and cardiovascular disease.

As further evidence of LRRC8 channel inhibition as a mechanism of action for anti-platelet activity, we tested dicoumarol, a newly identified VRAC inhibitor ^41^ that is structurally distinct from SN-40X compounds. As a 4-hydroxycoumarin anticoagulant, dicoumarol inhibits vitamin K-dependent clotting factor synthesis (II, VII, IX and X), resulting in impaired coagulation due to reduced clotting cascade efficacy ^74^. Based on the recent study by Chu *et al.* demonstrating dicoumarol-mediated blockade of LRRC8-mediated ATP release in HeLa cells and microglia ^41^, and our findings linking ATP efflux through LRRC8 channels in platelet activation, we speculated that dicoumarol would also exhibit anti-platelet activity. Our results suggest that, alongside dicoumarol’s known effects on vitamin K-dependent clotting factors, it also exhibits direct anti-platelet activity via inhibition of LRRC8 channel-mediated ATP efflux.

In summary, we identify a genetic signal for a modulatory role for LRRC8 proteins in platelet function in humans and validated this using platelet-targeted LRRC8A KO mice. We demonstrate that the platelet LRRC8 complex forms a novel mechanoresponsive ATP and ADP release channel that contributes to arterial thrombosis. Moreover, suppression of platelet function using small-molecule LRRC8 channel inhibitors highlights the opportunity to develop LRRC8 targeted anti-thrombotic agents.

## Methods

### Animals

All studies involving animal experimentation were performed in accordance with the recommendations in the Guide for the Care and Use of Laboratory Animals of the National Institutes of Health. All animals were handled according to the approved institutional animal care and use committee (IACUC) protocols of Washington University in St. Louis. C57BL/6N and C57BL/6J mice used in this study were obtained from Charles River Laboratories (Wilmington, MA, USA) and the Jackson Laboratory (Bar Harbor, ME, USA), respectively. Megakaryocyte (MK)-specific LRRC8A-knockout mice were created by crossing *Pf4*-Cre (Strain #008535, Jackson Laboratory) mice with the *Lrrc8a^fl/fl^* (*Swell1^fl/fl^*) mouse ^27^. All mice were housed in a 12 h light/dark cycle in a temperature and humidity-controlled room, with *ad libitum* access to standard chow and water.

### Bioinformatics

For genome-wide association studies (GWAS), the NHGARI-EBI catalog of eligible published human genome-wide association studies (https://www.ebi.ac.uk/gwas/home; accessed 9^th^ February 2022) was utilized to identify SNPs associated with mean platelet volume using a pre-specified p-value cutoff of 10^-8^. For phenome-wide association studies (PheWAS), the IEU Open GWAS project (https://gwas.mrcieu.ac.uk/; accessed 3^rd^ March 2022) was utilized to identify associations between each of the SNP variants identified in the GWAS search.

The Plateletomics Interactive Results Tool version 1.0 was used to search the Plateletomics database (plateletomics.com; accessed 5^th^ March 2022) of 5911 uniquely mapped mRNAs derived from the leukocyte-depleted RNA of 154 human subjects ^50^ for relative expression of LRRC8 subunits A – D. Similarly, this Plateletomics Interactive Results Tool was used to determine whether the extent of *in vitro* platelet aggregation (0 – 100%) by four established platelet agonists was associated with mRNA levels of LRRC8A – D. Additionally, the database found in Supplementary Table S2A from the 2014 publication by Simon *et al.* ^50^ (accessed 5^th^ March 2022) was used to determine the relative expression of transcripts for subunits LRRC8B, C and D, as compared against those for LRRC8A.

### Murine platelet isolation

Mice were anesthetized using either inhalational isoflurane (4% induction, 2.5 – 3% for maintenance) or intraperitoneal injection of a ketamine / xylazine cocktail (100 mg/kg and 10 mg/kg, respectively) in saline. Anesthetic depth was confirmed by testing reflexes in response to both tail tip and toe tip pinching. Blood was isolated from the inferior vena cava using a 26½ gauge needle into 1 mL syringes containing 100 µL acid citrate dextrose (ACD) solution (#C3821, Sigma-Aldrich, St. Louis, MO, USA) with subsequent centrifugation (300 x g; RT; 20 min; acceleration and brake both set to 0, swinging-bucket rotor). Supernatants and one-third of RBCs were then collected and mixed with 0.5 µM prostaglandin E1 (#13010, Cayman Chemical, Ann Arbor, MI, USA) and centrifuged (700 x g; RT; 5 min; acceleration = 1, brake = 0, swinging-bucket rotor). Resulting supernatants were then collected and further centrifuged (750 x g; RT; 5 min; fixed-angle rotor) to pellet platelets. Platelets were washed once by resuspension in HEPES-Tyrode’s buffer (HTB; 136 mM NaCl, 12 mM NaHCO_3_, 2.6 mM KCl, 2.5 mM HEPES, 5.5 mM glucose, CaCl_2_*, 1 mM MgCl_2_; made in molecular grade water) containing 10% ACD prior to further centrifugation and resuspension in 200 µL HTB. Platelet counts and MPVs were obtained using the VETSCAN^®^ HM5 Hematology Analyzer (Zoetis, Parsippany, NJ, USA). Platelets were then diluted to the desired concentration, which were assay dependent. * The amount of CaCl_2_ present in HTB at the time of platelet resuspension was dependent on which assay was being performed. In most assays, further CaCl_2_ was later added so that the final concentration of CaCl_2_ in each assay was 1 mM. Exact details of platelet concentration and CaCl_2_ are found under each assay’s specific methods.

### Human platelet isolation

Human whole blood was collected in accordance with the Institutional Review Board of Washington University, St. Louis from consenting healthy adults. Whole blood was collected by venipuncture into acid-citrate-dextrose (ACD) vacutainers (BD Biosciences, Franklin Lakes, NJ, USA). Blood was supplemented with 0.5 µM prostaglandin E1 (Cayman Chemical). Platelet-rich plasma (PRP) was fractionated from red blood cells (RBCs) by centrifugation (300 x g; RT; 20 min; acceleration and brake both set to 0). PRP was then supplemented with 0.5 µM prostaglandin E1 (Cayman Chemical), followed by further centrifugation (800 x g; RT; 20 min; acceleration set to 1, brake set to 0) to pellet platelets. Resulting supernatants were aspirated, followed by washing of platelet pellets in 10 mL of wash buffer (10% ACD in HTB). Washed pellets were then resuspended in 200 µL HTB (136 mM NaCl, 12 mM NaHCO_3_, 2.6 mM KCl, 2.5 mM HEPES, 5.5 mM glucose, CaCl_2_*, 1 mM MgCl_2_; made in molecular grade water) and platelet counts were obtained using the VETSCAN^®^ HM5 Hematology Analyzer (Zoetis). Platelets were then diluted to the desired concentration, which were assay dependent. * The amount of CaCl_2_ present in HTB at the time of platelet resuspension was dependent on which assay was being performed. In most assays, further CaCl_2_ was later added so that the final concentration of CaCl_2_ in each assay was 1 mM. Exact details of platelet concentration and CaCl_2_ are found under each assay’s specific methods.

### Western blot

Samples were lysed in ice cold radioimmunoprecipitation assay (RIPA) buffer containing 1:10 PhosSTOP™ (#4906845001, Roche Diagnostics, Indianapolis, IN, USA) and 1:20 cOmplete™ ULTRA Protease Inhibitor Cocktail (#06538304001, Roche Diagnostics) and were stored at -80°C overnight. Samples were then thawed and further lysed by sonication (20% amplitude with a 20 sec cycle interval, twice), followed by centrifugation (13,000 rpm; 20 min; RT). Resulting supernatants were collected and protein quantification was then performed using the DC™ Protein Assay Kit I (#5000111, Bio-Rad, Hercules, CA, USA). Samples were prepared by addition of 4X Laemmli Sample Buffer (#1610747, Bio-Rad) containing 1:10 β-mercaptoethanol and subsequent incubation at 95°C for 5 min prior to cooling on ice. SDS-PAGE was performed by loading 20 µg of each sample into 4 – 15% Mini-PROTEAN^®^ TGX™ Precast Protein Gels (#4561083, Bio-Rad) which were ran in tris/glycine/SDS running buffer (#1610772, Bio-Rad) at 100 V for 1.5 – 2h. Gels were transferred onto Immun-Blot PVDF Membranes (#1620177, Bio-Rad) in tris/glycine buffer (#1610771, Bio-Rad) for at 100 V for 1.5 h at 4°C. Membranes were then blocked in 5% bovine serum albumin (#A-420-10, GoldBio, St. Louis, MO, USA), or milk, in tris-buffered saline with Tween (TBS-T; 0.2 M Tris, 1.37 M NaCl, 0.2% Tween-20, pH 7.4) at room temperature for 1 h. Membranes were subsequently incubated with rabbit polyclonal anti-LRRC8A against the epitope QRTKSRIEQGIVDRSE ^29^ (Pacific Immunology, Ramona, CA, USA), anti-LRRC8B*, anti-LRRC8C (#21601-1-AP, Proteintech, Rosemont, IL, USA), anti-LRRC8D*, anti-LRRC8E*, anti-AKT1 (#2938, Cell Signaling Technology, Danvers, MA, USA), anti-pAKT1^Ser473^ (#9018, Cell Signaling Technology), anti-AKT2 (#3063, Cell Signaling Technology), anti-pAKT2^Ser474^ (#8599, Cell Signaling Technology), anti-P-selectin (#AF737, R&D Systems, Minneapolis, MN, USA), anti-GSK-3β (#12456, Cell Signaling Technology), anti-pGSK-3β^Ser9^ (#9323, Cell Signaling Technology), anti-Integrin β3 (#sc-6627, Santa Cruz Biotechnology, Dallas, TX, USA), Anti-pPI3K p85^Tyr458^ / p55^Tyr199^ (#4228, Cell Signaling Technology), anti-β-Actin (#8457, Cell Signaling Technology), or anti-GAPDH (#5174, Cell Signaling Technology) at a concentration of 1:1000 in 1% BSA or milk overnight at 4°C (* Indicates antibodies which were kindly provided by T.J. Jentsch, previously validated in KO cell lines ^75^). Membranes were washed three times in TBS-T prior to incubation with goat anti-rabbit IgG (H+L)-Horseradish Peroxidase Conjugate (#1706515, Bio-Rad) or donkey anti-Goat IgG (H+L)-Horseradish Peroxidase Conjugate (Invitrogen #A15999, Thermo Fisher Scientific, Waltham, MA, USA) at a concentration of 1:5000 in 1% BSA or milk at room temperature for 1 h prior to further washing. Chemiluminescence was achieved using Clarity™ Western ECL Substrate, (#1705060, Bio-Rad) or SuperSignal™ West Dura Extended Duration Substrate (#34075, Thermo Fisher Scientific) with imaging performed on the ChemiDoc Imaging System (Bio-Rad). Images were analyzed and blots quantified using Fiji (Version 2.14.0/1.54f ^76^).

### Murine whole blood isolation and complete blood counts

For CBC measurements in whole blood, mice were anesthetized using inhalational isoflurane (4% induction) and whole blood was obtained from the retro-orbital plexus using heparinized capillary tubes. Complete blood counts (CBCs) were evaluated using the Heska HT5 analyzer (Heska, Loveland, CO, USA).

### Murine megakaryocyte isolation

Mice were anesthetized by isoflurane (4% induction, 2.5 – 3% for maintenance) and, after cervical dislocation, femur and tibia bone were dissected out and kept in freshly prepared CATCH buffer (10X calcium-magnesium free Hank’s solution (CMF-HBSS) 100 mL; Milli-Q water 850 mL; adenosine 376 mg; theophylline 494 mg; sodium citrate 3.8 g; adjusted to 1L final volume and pH 7.4). Dissected femur and tibia were cleared from adherent muscle tissue and a cut-hole was made on both sides of the bone so that bone marrow (BM) could be flushed out via use of a 10 mL syringe filled CATCH buffer, with BM being collected in 50 mL conical tube. Subsequent centrifugation (200 x g; 10 min; RT) was performed to collect intact bone marrow pellets, followed by their resuspension in 1 mL of CATCH buffer. Resuspended BM cells were then layered gently onto a freshly prepared BSA stock-CATCH buffer gradient tube (bottom: BSA:CATCH, 5 : 16; middle: 3.3 : 10.6; top: 1.6 : 5.3 (V/V)) and allowed to separate for 30 min based on their density gradient (BSA stock solution (25 mL): Milli-Q water: 18 mL, 10X CMF-HBSS 1.9 mL; adenosine 15 mg; theophylline 16.05 mg; BSA 7.25 g; adjusted to 25 mL and pH 7.4). Subsequently, the top two layers of BSA gradient solution were removed carefully and remaining MK-rich bottom layer was resuspended with CATCH buffer and centrifuged (200 x g; 10 min; RT). Resulting supernatants were aspirated, and pellets of MKs were resuspended in CATCH buffer.

### Electrophysiology

All experiments were conducted at room temperature using either an Axopatch 200B amplifier or a MultiClamp 700B amplifier paired to a Digidata 1550 digitizer and both used pClamp 10.4 (Molecular Devices, San Jose, CA, USA) software. For standard VRAC whole-cell recordings, the intracellular solution contained (in mM): 120 L-aspartic acid, 20 CsCl, 1 MgCl_2_, 5 EGTA, 10 HEPES, 5 MgATP, 120 CsOH, 0.1 GTP, pH 7.2 with CsOH. For measuring ATP and ADP currents, two sets of intracellular solutions were prepared at concentrations of 50 mM and 1 mM. For the ATP solutions, the 50 mM intracellular solution contained (mM): 50 Na_2_ATP^2-^, 10 HEPES, 150 mannitol, pH 7.2 with NaOH, while the 1 mM solution contained (mM): 1 Na_2_ATP^2-^, 10 HEPES, 150 mannitol, 120 Aspartic Acid, 120 CsOH, pH 7.2 with NaOH. For the ADP solutions, the 50 mM intracellular solution contained (mM): 50 Na_2_ADP, 1 Na_2_ATP, 10 HEPES, 150 Mannitol, pH 7.2 with NaOH, while the 1 mM solution contained (mM): 1 Na_2_ADP, 1 Na_2_ATP, 10 HEPES, 150 Mannitol, 120 Aspartic Acid, 120 CsOH, pH 7.2 with NaOH. The isotonic extracellular solution contained (in mM): 90 NaCl, 2 CsCl, 1 MgCl_2_, 1 CaCl_2_, 10 HEPES, 110 mannitol, pH 7.4 with NaOH (300 mOsm/kg). For hypotonic swelling (210 mOsm/kg), the extracellular solution contained the same composition above but with 10 mM mannitol instead of 110 mM. Swell-activated VRAC was elicited by perfusing cells with hypotonic solution (210 mOsm/kg). Osmolarities were measured using the Wescor Vapro 5520 Osmometer (ELITechGroup Inc, Logan, UT, USA). The patch electrodes were prepared from borosilicate glass capillaries (WPI) and had a resistance of 2.5 – 4.8 MΩ when filled with pipette solution. For perforated patch recordings, the intracellular solution was as above but without ATP and GTP, and contained 360 µg/mL Amphotericin B (#A2942, Sigma-Aldrich). The holding potential was 0 mV. Voltage ramps from −100 to +100 mV (at 0.4 mV/ms) were applied every 4 sec for standard VRAC measurements, and -140 mV to -80 mV every 4 sec for ATP and ADP currents. Sampling interval was 100 µs and filtered at 10 KHz. For perforated-patch recordings, cells with a membrane resistance below GΩ or access resistance above 20 MΩ were discarded.

### Microfluidic platelet adhesion

Platelet adhesion to collagen under physiological shear conditions was performed using flow-based microfluidics. Type I collagen (#385, Chrono-Log Corporation, Havertown, PA, USA) was patterned for 2 h onto a glass slide. A polydimethylsiloxane microfluidic flow chamber was vacuum sealed to the glass slide and flow channels were blocked with 2% BSA for 1 h. Murine whole blood was collected into heparin anticoagulant via the IVC and platelets were fluorescently labelled with Alexa Fluor^®^ 488 anti-mouse CD41 antibody (#133907, BioLegend, San Diego, CA, USA). Whole blood was perfused through the channels at a wall shear rate of 650 s^−1^ for 5 min using a Harvard syringe pump and platelet adhesion was captured in real time fluorescence using an Olympus IX83 microscope (Olympus, Center Valley, PA, USA). Captured images and platelet surface area was analyzed using cellSens software (Olympus).

### Flow cytometry

In the study investigating murine platelet activation in response to different agonists, platelets were isolated from *Lrrc8a^fl/fl^* (WT) and *Pf4-Cre;Lrrc8a^fl/fl^* (cKO) mice were resuspended and normalized to 300 x 10^6^ platelets / µL in HTB containing 1 mM CaCl_2_. Samples in 10 µL aliquots were then stimulated with 1 µL of thrombin (#T7009, Sigma-Aldrich, 0.03 U/mL final concentration), CRP (#CRP-XL, Cambcol Laboratories, Ely, Cambridgeshire, UK, 0.2 µg/mL final concentration), or A23187 (#C7522, Sigma-Aldrich, 0.5 µM final concentration) for 5 min at 37°C, followed by incubation with 1 µL of an equal mix of FITC-labeled rat anti-mouse P-selectin (CD62P) (#M130-1, Emfret Analytics, Würzburg, Germany) and PE-labeled rat anti-mouse integrin αIIbβ3 (GPIIbIIIa, CD41/61) (#M023-2, Emfret Analytics) in the dark at room temperature for 20 min. Samples were subsequently fixed in 300 µL of 1% paraformaldehyde (PFA) in phosphate-buffered saline (PBS). P-selectin exposure and JonA binding (to activated αIIbβ3 integrin) were analyzed using a flow cytometer (CytoFLEX, Beckman Coulter, Indianapolis, IN, USA).

In the study investigating the small-molecule LRRC8 complex inhibitor SN-89B on agonist-induced human and murine platelet activation, both human and mouse platelets were resuspended and normalized to 300 x 10^6^ platelets / µL in HTB containing 200 µM CaCl_2_ supplemented with either vehicle (0.02% DMSO) or SN-89B (10 µM). Prior to activation, 10 µL of each sample was gently mixed with 10 mM CaCl_2_ such that the final concentration was 1 mM. Murine samples were then stimulated with PAR4-AP (400 µM final concentration), whereas human samples were stimulated with a cocktail of PAR1-AP (10 µM final concentration) and PAR4-AP (100 µM final concentration), for 5 min at 37°C. Murine samples were then incubated in the dark at room temperature with 1.2 µL of an equal mix of FITC-labeled rat anti-mouse P-selectin (CD62P) (Emfret Analytics) and PE-labeled rat anti-mouse integrin αIIbβ3 (Emfret Analytics).

Human samples were incubated in the dark at room temperature with 1.2 µL of an equal mix of BD Pharmingen™ PE Mouse Anti-Human CD62P (BD Biosciences) and BD™ FITC Mouse Anti-Human PAC-1 (#340507, BD Biosciences). All samples were subsequently fixed in 300 µL of 1% PFA in PBS. P-selectin exposure (all), JonA binding (murine only) and PAC-1 binding (human only) to activated αIIbβ3 integrin, were then analyzed using a flow cytometer (CytoFLEX, Beckman Coulter).

### Human iPSC-derived megakaryocytes

Experiments using CD34^+^-derived megakaryocytes were approved by the Institutional Review Boards of both the University of Utah and Washington University in St. Louis. CD34^+^ hematopoietic progenitors were isolated from human umbilical cords and differentiated into MKs, followed by transfection, as previously described by Montenont *et al.*^52^. On day 5, CD34^+^ cells were transfected using predesigned Alt-R™ CRISPR-Cas9 guide (g)RNA (Integrated DNA Technologies, Coralville, IA, USA) targeting LRRC8A (ACCAAGTCACGGATCGAGCA), or control gRNA. On day 13, 100 µL of MKs resuspended at 10^6^ per mL were stained with BD Pharmingen™ APC Mouse Anti-Human CD41a (#559777, BD Biosciences), BD™ FITC Mouse Anti-Human PAC-1 (#340507, BD Biosciences) and BD Pharmingen™ PE Mouse Anti-Human CD62P (#555524, BD Biosciences), and were subsequently activated for 15 min at 37 °C by treatment with 0.1, 0.25 or 0.5 U/mL thrombin (#T7009, Sigma-Aldrich). MKs were subsequently fixed with 2% paraformaldehyde and activation measured using the CytoFLEX flow cytometer apparatus (Beckman Coulter). Normalized mean florescence intensity (MFI) was used to determine the number of cells bound to P-selectin and PAC-1. MFI of activated LRRC8A KO cells were compared to the MFI of activated negative control cells derived from the same cord.

### Light transmission aggregometry

Platelet aggregation and ATP release were measured using the CHRONO-LOG^®^ Model 700 apparatus (Chrono-Log Corporation). In all instances, samples were incubated for a minimum of 5 min at 37°C prior to aggregometry. All samples were added in 250 µL volumes into glass cuvettes containing magnetic stir bars, with certain samples also having 10 µL of CHRONO-LUME^®^ luminescent reagent (#395, Chrono-Log Corporation) added to measure ATP release. All samples were incubated at 37°C and stirred at 1000 rpm during aggregometry.

For the study investigating the effects of thrombin, CRP, and U46619 on murine platelet aggregation, platelets isolated from *Lrrc8a^fl/fl^* (WT) and *Pf4-Cre;Lrrc8a^fl/fl^* (cKO) mice were resuspended and normalized to 300 x 10^6^ platelets / µL in HTB containing 1 mM CaCl_2_. After determining a stable baseline in each sample, ADP (10 – 20 µM, #21121, Cayman Chemical) with fibrinogen (30 µg/mL, #341576, Sigma-Aldrich); CRP (0.05 – 0.15 µg/mL, #CRP-XL, CambCol Laboratories); U46619 (0.5 – 0.8 µM, #16450, Cayman Chemical) or thrombin (0.02 – 0.05 U/mL, #T7009, Sigma-Aldrich) were added, followed by measurements of platelet aggregation and ATP release for up to 10 min.

For the study investigating the effects of hypotonic swelling on platelet aggregation in the presence or absence of SN-401, human platelets were resuspended and normalized to 600 x 10^6^ platelets / µL in HTB containing 200 µM CaCl_2_. HTB was supplemented with either vehicle (0.02% DMSO) or SN-401 (10 µM). After determining a stable baseline in each sample, 125 µL of pre-heated (37°C) molecular grade water containing either vehicle (0.02% DMSO) or SN-401 (10 µM) were added, reducing HTB osmolarity to ∼ 180 mOsm (‘hypotonic’). Platelet aggregation was then measured for 10 min.

For the study investigating the effects of hypotonic swelling on platelet aggregation in the presence or absence of TNP-ATP or MRS2179, human platelets were resuspended and normalized to 600 x 10^6^ platelets / µL in HTB containing 200 µM CaCl_2_. HTB was supplemented with TNP-ATP (850nM) (#20902, Cayman Chemical) or MRS2179 (700 nM) (#0900, Tocris Bioscience, Bristol, UK). After determining a stable baseline in each sample, 125 µL of pre-heated (37°C) molecular grade water with or without TNP-ATP (850nM) or MRS2179 (700 nM) were added, reducing HTB osmolarity to ∼ 180 mOsm (‘hypotonic’). Platelet aggregation was then measured for 10 min.

For studies investigating the effects of hypotonic swelling on platelet aggregation in the presence or absence of NF449 or ticagrelor, human platelets were resuspended and normalized to 300 x 10^6^ platelets / µL in HTB containing 200 µM CaCl_2_. HTB was supplemented with either vehicle (0.02% DMSO), NF449 (500 nM or 50 µM) (#13324, Cayman Chemical) or ticagrelor (85 nM) (#SML2482, Sigma-Aldrich). After determining a stable baseline in each sample, 125 µL of pre-heated (37°C) molecular grade water containing either vehicle (0.02% DMSO), NF449 (500 nM or 50 µM), or ticagrelor (85 nM) were added, reducing HTB osmolarity to ∼ 180 mOsm (‘hypotonic’). Platelet aggregation was then measured for 10 min.

For studies investigating the effects of small-molecule LRRC8 channel inhibitors on platelet aggregation and ATP release, human and mouse platelets were resuspended and normalized to 300 x 10^6^ platelets / µL in HTB containing 200 µM CaCl_2_. HTB was supplemented with vehicle (0.02% DMSO), SN-401, SN-406, SN-407, or SN-89B (10 µM). For studies in which the half-maximal inhibitory concentration (IC_50_) of SN-406 (using murine platelets) and SN-89B (using human platelets) were calculated, a range of concentrations were used. Murine platelets were subsequently activated with thrombin (#386, Chrono-Log Corporation, 0.02 U/mL final concentration) or PAR4-AP (400 µM final concentration). Human platelets were activated with a cocktail of PAR1-AP (10 µM final concentration) and PAR4-AP (100 µM final concentration). All agonists were supplemented with CaCl_2_ such that the final concentration of Ca^2+^ in each sample was 1 mM. In instances where inhibition (%) was calculated, the mean aggregation of platelets at each tested drug concentration was compared against the maximal response of platelets treated with vehicle control only.

For the study investigating the inhibitory effects of dicoumarol on platelet aggregation, platelets isolated from WT C57BL/6J mice were normalized to 300 x 10^6^ platelets / µL in HTB supplemented with vehicle (0.1% DMSO) or dicoumarol (20 µM, #204120050, Thermo Fisher Scientific) prior to aggregometry. After determining a stable baseline in each sample, aggregation was induced by addition of PAR4-AP (400 µM final concentration). Platelet aggregation was measured for 20 min, with addition of 40 µM ATP (#387, Chrono-Log Corporation) at 10 min. PAR4-AP was supplemented with CaCl_2_ such that the final concentration of Ca^2+^ in each sample was 1 mM.

### Calcium flux assay

Aliquots of 100 µL platelets, normalized to either 200 x 10^6^ platelets / µL (murine) or 300 x 10^6^ platelets / µL (human) in HTB containing 200 µM CaCl_2_, were mixed with 50 µL of Component A from the Fura-2 QBT Calcium Kit (#R6193, Molecular Devices) and incubated in the dark at room temperature for 30 min. For drug studies, HTB and Component A were supplemented with vehicle or compounds. After incubation, 27 µL of each sample was added to clear bottom 384-well plates. Using the FlexStation^®^ 3 Multi-Mode Microplate Reader (Molecular Devices), samples were incubated at 37°C for 5 min prior to initializing a program set to read fluorescence at 340 nm and 380 nm every 5 sec for a total of 10 min. For the study in which calcium transients were evaluated between *Lrrc8a^fl/fl^* (WT) littermate control and *Pf4-Cre;Lrrc8a^fl/fl^* (cKO) mice, 3 µL of thrombin (#386, Chrono-Log Corporation, 0.02 U/mL final concentration) was added at 60 sec. For drug studies using murine platelets, either thrombin, as previously described, or 3 µL of PAR4-AP (400 µM final concentration), was added at 60 sec. For drug studies using human platelets, a cocktail of PAR1-AP (10 µM final concentration) and PAR4-AP (100 µM final concentration) was added at 60 sec. In all instances, agonists were supplemented with CaCl_2_ such that the final concentration of calcium in each sample was 1 mM.

### Signaling studies

For signaling studies utilizing platelets, platelets isolated from *Lrrc8a^fl/fl^* (WT) and *Pf4-Cre;Lrrc8a^fl/fl^* (cKO) mice were resuspended and normalized to 300 x 10^6^ platelets / µL in HTB containing 200 µM CaCl_2_. Light transmission aggregometry was performed as previously described, with 60 µL RIPA buffer added at 5 min to lyse each sample. Western blots were then performed as previously described.

For signaling studies utilizing MEG-01 cells, MEG-01 cells were maintained and transduced as described under ‘Cell culture’. Cells were stimulated with 2.5 U/mL thrombin for 15 min at 37°C, followed by addition of RIPA buffer to lyse cells. Western blots were then performed as previously described.

### Intravital microscopy

Intravital microscopy was performed in a mouse model of laser-induced cremaster arteriolar injury as described previously ^77^. *Lrrc8a^fl/fl^* (WT) littermate control and *Pf4-Cre;Lrrc8a^fl/fl^* (cKO) male mice (8 – 10 weeks old) were anesthetized by intraperitoneal injection of a ketamine/xylazine cocktail (100 mg/kg and 10 mg/kg, respectively) in saline. The cremaster muscle was exteriorized and superfused with 37 °C bicarbonate-buffered saline throughout the experiment. Arteriolar wall injury was induced by laser ablation (Ablate!™, Intelligent Imaging Innovations, Denver, CO, USA). Multiple thrombi were generated in at least 3 – 4 different arterioles of one mouse, with new thrombi formed upstream of earlier thrombi to minimize any contribution from thrombi generated earlier in the mouse. Platelets were visualized by injection of DyLight 649-conjugated anti-CD42c antibody (#X649, Emfret Analytics, 0.2 µg/g B.W.). Fluorescence and bright-field images were captured in 6 – 10 different cremaster arterioles with a diameter of 30 – 45 µm in each mouse and recorded using a Zeiss Axio examiner Z1 microscope system with a high-power LED light source (X-Cite XLED1 5-Channel Light Source (390 – 415 nm, 450 – 495 nm, 505 – 545 nm, 540 – 600 nm, and 615 – 655 nm, Excelitas Technologies, Waltham, MA, USA). Images were collected with a high-speed, high-resolution camera (2304 x 2304 pixel format, ORCA-Fusion BT sCMOS, Hamamatsu, Shizuoka, Japan). Time 0 was set to when image capture began on each vessel. Data were analyzed using Slidebook (version 6.0, Intelligent Imaging Innovations). Due to the significant variation between vessels in the same mouse, the result from each vessel was counted as an individual value. Representative images were chosen that most closely resemble the mean values in the quantifications. Experiments were performed in a single-blind fashion in which the investigators did not know the identity of the sample or mouse.

### FeCl_3_-induced carotid arterial thrombosis

For the study which investigated thrombosis in *Lrrc8a^fl/fl^* (WT) littermate control and *Pf4-Cre;Lrrc8a^fl/fl^* (cKO) mice, the right carotid artery was isolated from surrounding tissues. A nanoprobe (0.5PSB, Transonic Systems, Ithaca, NY, USA) was hooked to the artery, and blood flow was monitored with a TS420 flowmeter (Transonic Systems). After stabilization, 1.2 µL of 7% FeCl_3_ (#908908, Sigma-Aldrich) was applied to a filter paper disc (2 x 2 mm), and the paper was placed on the top of the artery for 2 min. After removing the filter paper, blood flow was monitored continuously until 5 min after occlusion. The time to occlusion was measured as a difference in time between the removal of the filter paper and stable occlusion (no blood flow for 5 min).

For the study in which potential antithrombotic effects of small-molecule LRRC8 complex inhibitor SN-89B was evaluated, the procedure was the same as previously described, but used 5% FeCl_3_ (Sigma-Aldrich).

### Bleeding time assay

*Lrrc8a^fl/fl^* (WT) littermate control and *Pf4-Cre;Lrrc8a^fl/fl^*(cKO) mice were anesthetized via intraperitoneal injection of a ketamine/xylazine cocktail as previously described. Tail segments of 5 mm in length were amputated, followed by immersion of the tail into pre-warmed physiological saline. Bleeding was monitored for 15 min. Following cessation of bleeding, tails were monitored for an additional 5 min to ensure no rebleeding. In mice which failed to clot after 15 min, hemostasis was achieved using silver nitrate. Tubes of saline/blood were subsequently centrifuged (3900 rpm; RT; 10 min), with resulting supernatants being aspirated and pelleted RBCs were lysed in 2 mL of eBioscience™ 1X RBC Lysis Buffer (#00-4333-57, Thermo Fisher Scientific) for 60 min. Lysed RBCs were then centrifuged (12,000 rpm; RT; 20 min), with resulting supernatants collected for analysis of a hemoglobin concentration by measuring absorbance at 550 nm.

### Cell culture

MEG-01 megakaryoblasts (#CRL-2021, ATCC, Manassas, VA, USA) were maintained in RPMI 1640 medium (Gibco #11875085, Thermo Fisher Scientific) containing 10% fetal bovine serum (FBS, #S11550, R&D Systems) and 100 U/mL penicillin and 100 µg/ml streptomycin (Gibco #15140122, Thermo Fisher Scientific) at 37°C and 5% CO_2_. For LRRC8A KD studies, cells were transduced with either Ad-shLRRC8A-mCherry or Ad-shSCR-mCherry control for 72 h (MOI = 20) prior to either stimulation with 2.5 U/mL thrombin for 15 min and subsequent lysis as described under ‘Western blot’, or were seeded for 2 h at 37°C and 5% CO_2_ onto glass coverslips using 20 µg/mL human fibronectin (#354008, Corning, Corning, NY, USA) prior to patch clamp.

## Supporting information

Visual Abstract

## Author contributions

Conceptualization: R.S., D.J.L., J.D.T., A.K., R.M.

Formal Analysis: J.D.T., R.M., A.K., G.B., T.M.A.E., Y.Z., N.A., J.C., D.J.L. N.O.S., R.S.

Investigation: J.D.T., R.M., A.K., G.B., D.J.L., N.O.S., T.M.A.E., Y.Z., N.A., C.M., A.A., L.X., K.A., J.H., H.Z., T.K., A.L., A.B., V.K.N., N.M.L., Y.F.

Writing – Original Draft: R.S., J.D.T., and R.M.

Writing – Review & Editing: R.S., J.D.T., R.M., G.B., J.C., D.J.L.

Funding Acquisition: R.S., J.C., R.A.C., J.D.P., D.J.L.

Visualization: J.D.T., A.K., G.B., D.J.L.

Resources: R.S., J.C., R.A.C, J.D.P., D.J.L.

Supervision: R.S., J.C., R.A.C., J.D.P.

## Acknowledgements

We wish to thank the High-Throughput Screening Center (HTSC) at Washington University in St. Louis for use of their FlexStation^®^ 3 apparatus, and also Dr. Thomas Jenstch for kindly sharing LRRC8B, LRRC8D, and LRRC8E rabbit polyclonal antibodies. This work was supported by NIH NIDDK R01DK106009 (R.S.), R01DK126068 (R.S.), R01DK127080 (R.S.), NIH NHLBI R01HL168600 (R.S.), I01 BX005072 (R.S.), NIDDK R43 DK121598 (D.J.L.), R44 DK126600 (D.J.L.), NIH NHLBI R44 HL169181-01A1 (D.J.L.), NIH NHLBI R01HL146559 (J.C.), R01HL148280 (J.C.), R01HL28731 (J.C.), NIH NHLBI R01 HL139825 (J.D.P.), R61HL141794 (J.D.P.), NIH NHLBI R01HL160808 (R.A.C), R01HL163019 (R.A.C) and American Society for Hematology Graduate Student Award (A.A.).

