## Supplementary Figures for "LRRC8 complexes are adenosine nucleotide release channels regulating platelet activation and arterial thrombosis"

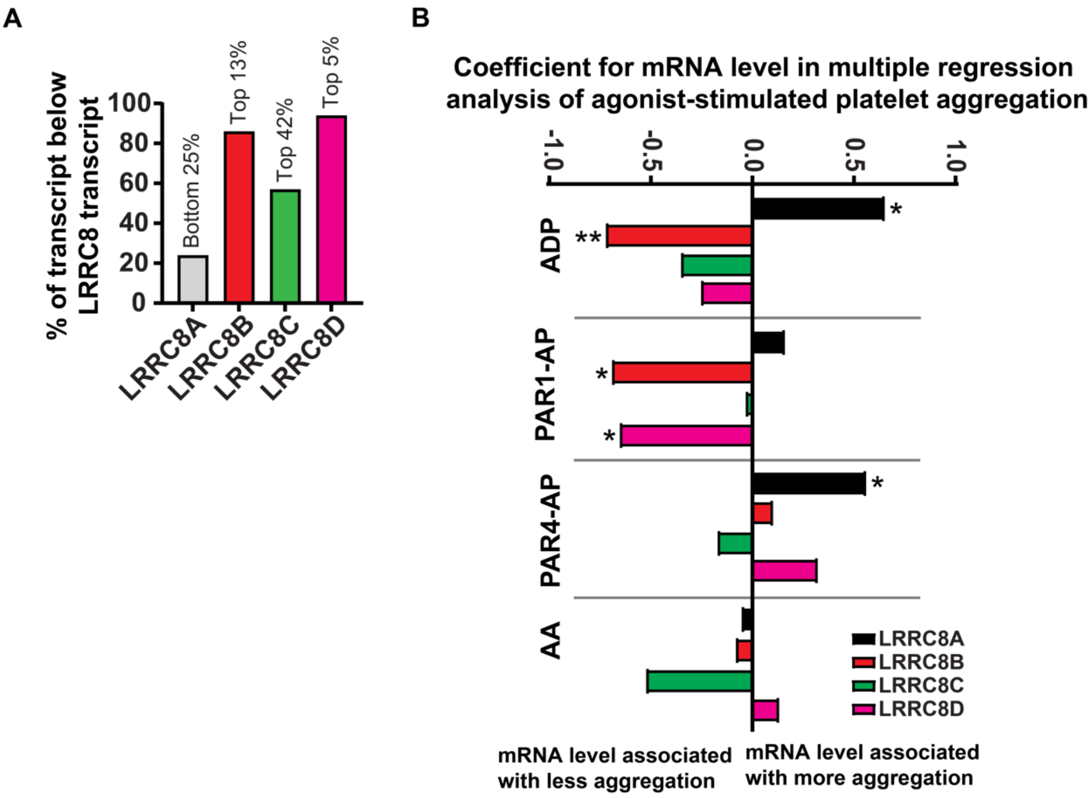


**Figure S1. Platelet express LRRC8 transcripts which are associated with altered agonist-induced aggregation. a** Ranked expression level of LRRC8 transcripts as compared against all other mRNA transcripts in human platelets (From Simon *et al.* [50] - Supplemental Table 2). **b** Coefficients for mRNA levels in multiple linear regression analysis of agonist response, where the dependent variable is agonist-aggregation response score for ADP, PAR1- and PAR4-activating peptides, and arachidonic acid (AA), and the independent variable is RNA level. *p* values in multiple linear regression model are derived from a T-test where T = coefficient/standard error (From Simon *et al.* [50] - Supplemental Table 3). *Abbreviations*: AA: arachidonic acid; ADP: adenosine diphosphate; AP; activating peptide; PAR: protease-activated receptor. * $p < 0.05$, ** $p < 0.01$.

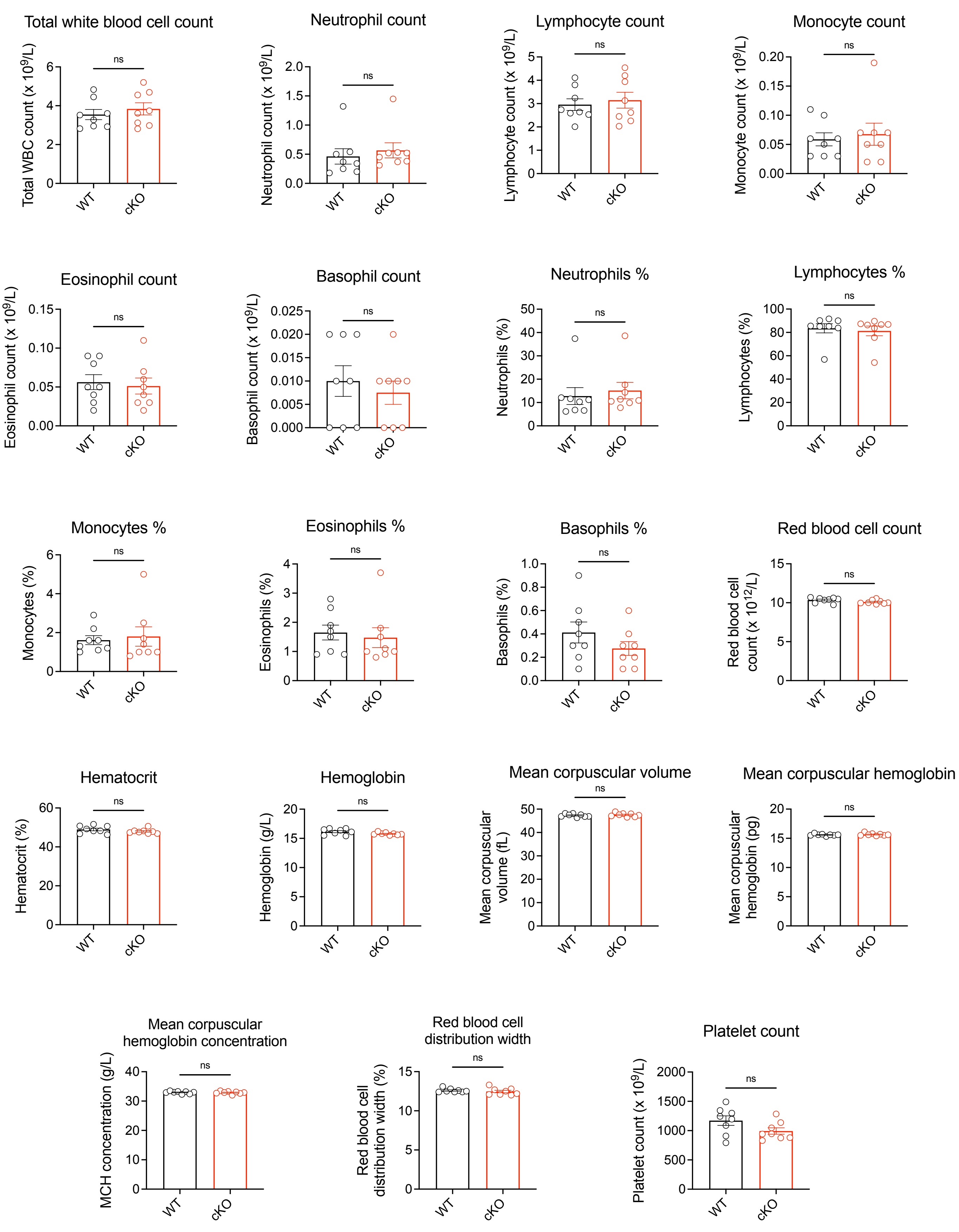


**Figure S2. Complete blood counts of platelet-specific LRRC8A KO mice.** Hematological parameters obtained by complete blood counts (CBCs) of whole blood isolated from *Lrrc8a*$^{fl/fl}$ (WT; n = 8) and *Pf4-Cre;Lrrc8a*$^{fl/fl}$ (cKO; n = 8) mice. Data are represented as mean ± S.E.M. Statistical significance was determined by unpaired T-test for all parameters. ns – not significant.

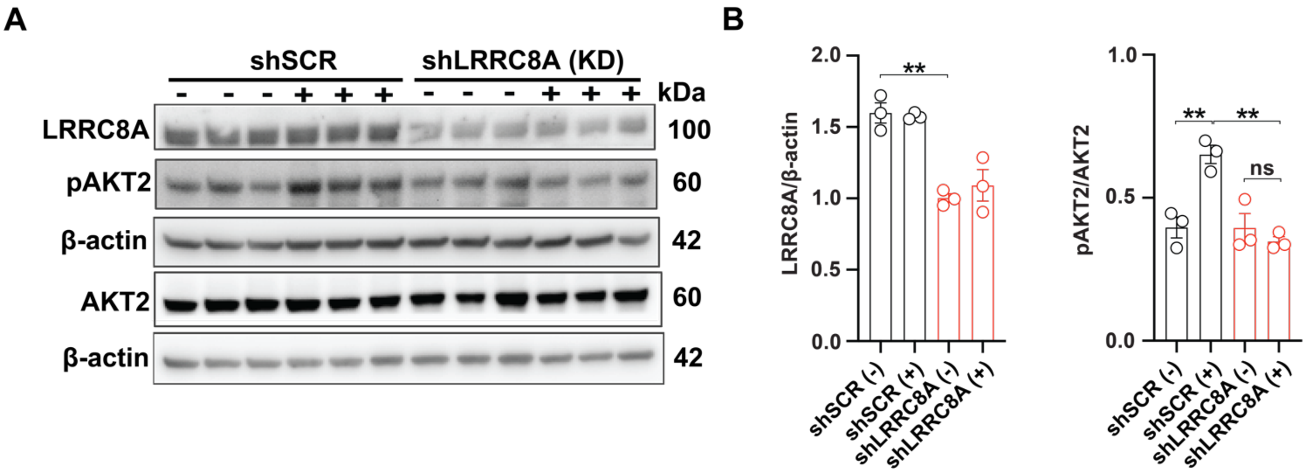


**Figure S3. LRRC8A knock-down impairs thrombin-induced AKT signaling in MEG-01 cells**. **a** Western blots detecting LRRC8A; AKT2; pAKT2$^{Ser474}$; and β-actin in MEG-01 cells transduced with adenoviral short hairpin control (shSCR) or one targeting LRRC8A (shLRRC8A; knock-down (KD)) with subsequent thrombin stimulation (2.5 U/mL, 15 min). **b** Densitometric quantification for LRRC8A / β-actin (n = 3) and pAKT2 / AKT2 (n = 3). Data are represented as mean ± S.E.M. Statistical significance was determined by unpaired T-test. ns – not significant; ** $p < 0.01$.

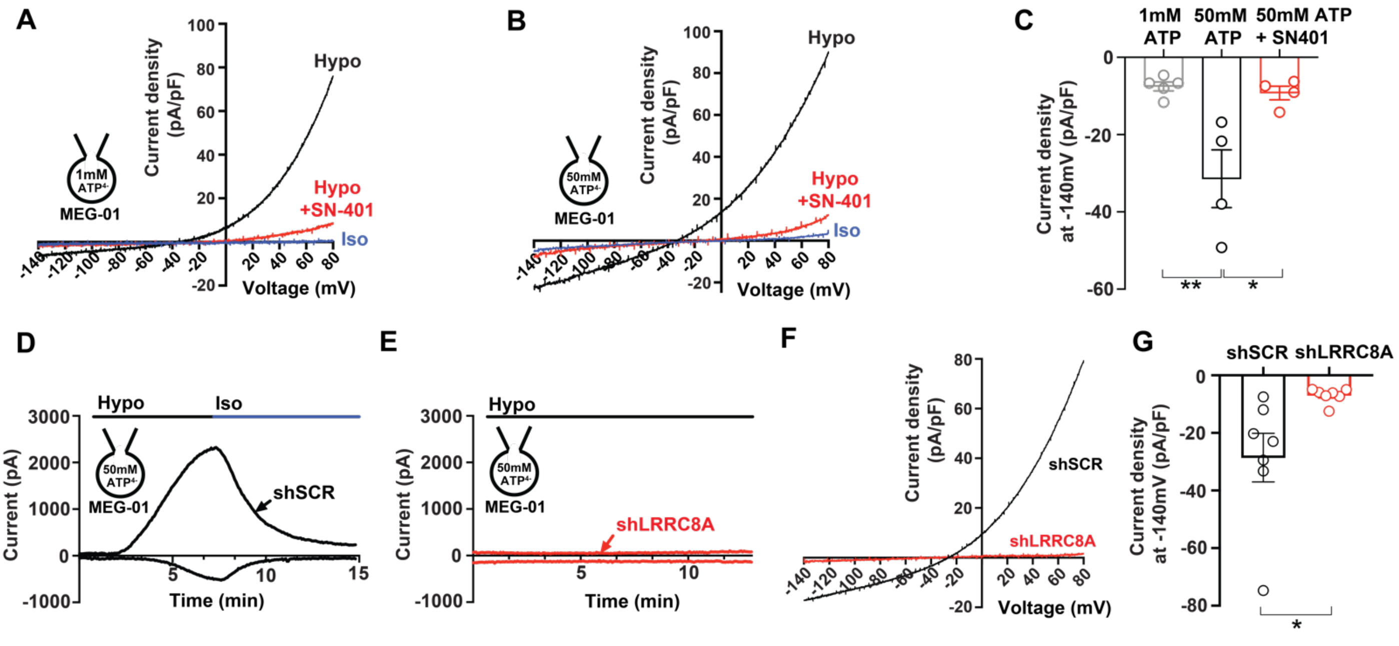


**Figure S4. MEG-01 LRRC8 channels permeate ATP. a, b** Current-voltage relationship of inward $I_{ATP}$ (ATP efflux) and outward VRAC in MEG-01 cells elicited from voltage ramps from -140 mV to +80 mV before and after hypotonic swelling with an intracellular ATP concentration of 1 mM (**a**) or 50 mM (**b**), followed by application of 10 μM SN-401 (DCPIB). The inward component of the current represents $I_{ATP}$ generated from ATP efflux**. c** Mean current densities of inward $I_{ATP}$ (ATP efflux) in MEG-01 cells at -140 mV after hypotonic swelling with an intracellular concentration of 1 mM ATP (n = 4), 50 mM ATP (n = 4), and 50 mM ATP + 10 μM SN-401 (n = 4). **d, e** Current-time relationship of inward $I_{ATP}$ and outward VRAC induced by hypotonic (210 mOsm) swelling in MEG-01 cells transduced with adenoviral short hairpin control (shSCR; **d**) or one targeting LRRC8A (shLRRC8A; **e**). **f** Current-voltage relationship of inward $I_{ATP}$ (ATP efflux) and outward VRAC during voltage ramps from -140 mV to +80 mV after hypotonic swelling in MEG-01 cells treated with either shSCR or shLRRC8A. **g** Mean current densities of inward $I_{ATP}$ (ATP efflux) in MEG-01 cells treated with shSCR (n = 7) or shLRRC8A (n = 7) at -140 mV following hypotonic swelling. Data are represented as mean ± S.E.M. Statistical significance was determined by unpaired t-test for **c** and **g**. * $p < 0.05$; ** $p < 0.01$.

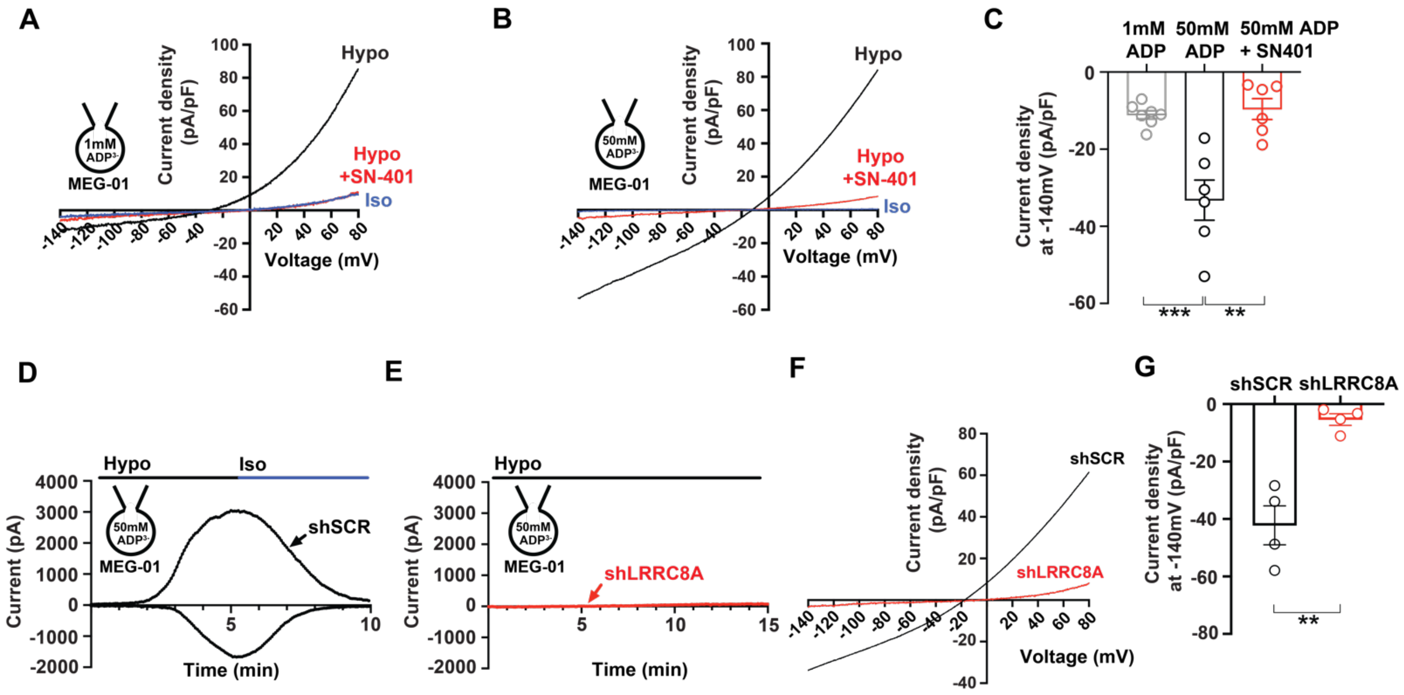


**Figure S5. MEG-01 LRRC8 channels permeate ADP. a, b** Current-voltage relationship of inward $I_{ADP}$ (ADP efflux) and outward VRAC in MEG-01 cells elicited from voltage ramps from -140 mV to +80 mV before and after hypotonic swelling with an intracellular ADP concentration of 1 mM (**a**) or 50 mM (**b**) followed by application of 10 µM SN-401 (DCPIB). The inward component of the current represents $I_{ADP}$ generated from ADP efflux**. c** Mean current densities of inward $I_{ADP}$ (ADP efflux) in MEG-01 cells at -140 mV after hypotonic swelling with an intracellular concentration of 1 mM ADP (n = 7), 50 mM ADP (n = 6), and 50 mM ADP + 10 µM SN-401 (n = 6). **d, e** Current-time relationship of inward $I_{ADP}$ and outward VRAC induced by hypotonic (210 mOsm) swelling in MEG-01 cells transduced with adenoviral short hairpin control (shSCR; **d**) or one targeting LRRC8A (shLRRC8A; **e**) for 72 h. **f** Current-voltage relationship of inward $I_{ADP}$ (ADP efflux) and outward VRAC during voltage ramps from -140 mV to +80 mV after hypotonic swelling in MEG-01 cells treated with either shSCR or shLRRC8A. **g** Mean current densities of inward $I_{ADP}$ (ADP efflux) in MEG-01 cells treated with shSCR (n = 4) or shLRRC8A (n = 4) at -140 mV following hypotonic swelling. Data are represented as mean ± S.E.M. Statistical significance was determined by unpaired T-test for **c** and **g**. ** $p < 0.01$; *** $p < 0.001$.

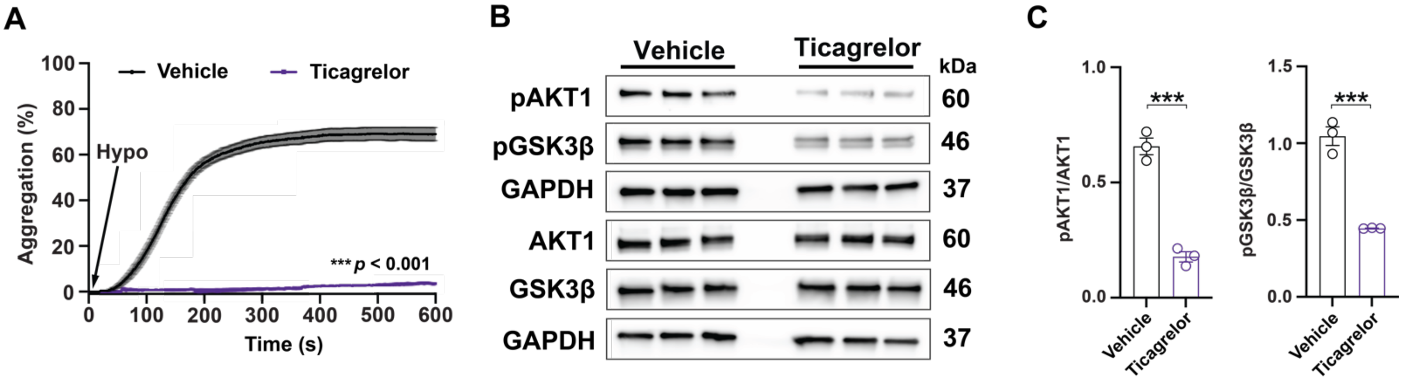


**Figure S6. P2Y$_{12}$ inhibitor ticagrelor inhibits hypotonic-induced aggregation and AKT1-GSK-3β signaling in murine platelets. a** Aggregometry of mouse platelets following the addition of molecular grade water to decrease HTB osmolarity from isotonic (270 mOsm) to hypotonic (180 mOsm) in the presence (n = 3 or absence (n = 3) of the P2Y$_{12}$ inhibitor ticagrelor (85 nM). **b** Western blots detecting pAKT1$^{Ser473}$; AKT1; pGSK-3β$^{Ser9}$; GSK-3β; and GAPDH in mouse platelets activated by decreasing HTB osmolarity in the presence (n = 3) or absence (n = 3) of the P2Y$_{12}$ inhibitor ticagrelor (85 nM), with densitometric quantification of pAKT1 / AKT1 and pGSK-3β / GSK-3β shown in **c**. Data are represented as mean ± S.E.M. Statistical significance was determined by two-way ANOVA for **a**, and by unpaired T-test for **c**. *** $p$ < 0.001.

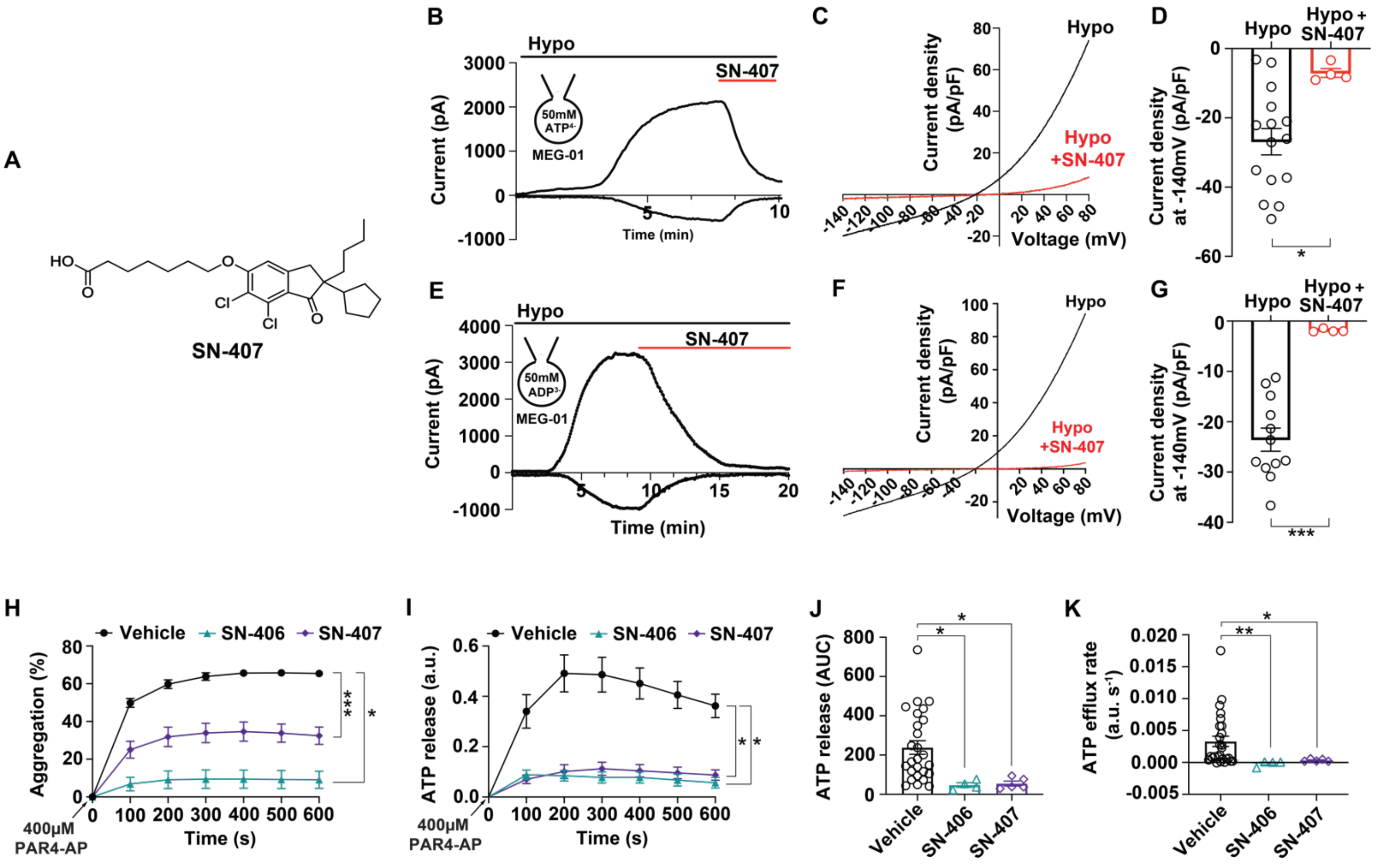


**Figure S7. LRRC8 channel small molecule inhibitors suppress agonist-induced aggregation and ATP release. a** Molecular structure of the small molecule LRRC8 channel complex modulator SN-407 – a derivative of SN-401 (DCPIB). **b** Current-time relationship of inward $I_{ATP}$ (ATP efflux) and outward VRAC induced by hypotonic (210 mOsm) swelling in MEG-01 cells and subsequent inhibition by application of 10 μM SN-407. **c, d** Current-voltage relationship of inward $I_{ATP}$ and outward VRAC during voltage ramps from -140 mV to +80 mV in MEG-01 cells after hypotonic swelling in the absence or presence of 10 μM SN-407, with mean current densities of inward $I_{ATP}$ at -140 mV shown in **d** (n = 4 – 15). **e** Current-time relationship of inward $I_{ADP}$ (ADP efflux) and outward VRAC induced by hypotonic (210 mOsm) swelling in MEG-01 cells and subsequent inhibition by application of 10 μM SN-407. **f, g** Current-voltage relationship of inward $I_{ADP}$ and outward VRAC during voltage ramps from -140 mV to +80 mV in MEG-01 cells after hypotonic swelling in the absence or presence of 10 μM SN-407, with mean current densities of inward $I_{ADP}$ at -140 mV shown in **g** (n = 4 – 12). **h – k** Aggregometry of platelets isolated from WT C57BL/6J mice stimulated with PAR4-AP (400 μM) in the presence of vehicle (0.02% DMSO; n = 29), SN-406 (10 μM; n = 5), or SN-407 (10 μM; n = 5), with concurrent ATP release (n = 4 – 26) shown in **i**, **j,** and **k**. Data are represented as mean ± S.E.M. Statistical significance was determined by unpaired T-test for **d**, **g** and **j**, Mann-Whitney for **k**, and two-way ANOVA for **h** and **i**. * $p < 0.05$; ** $p < 0.01$; *** $p < 0.001$.

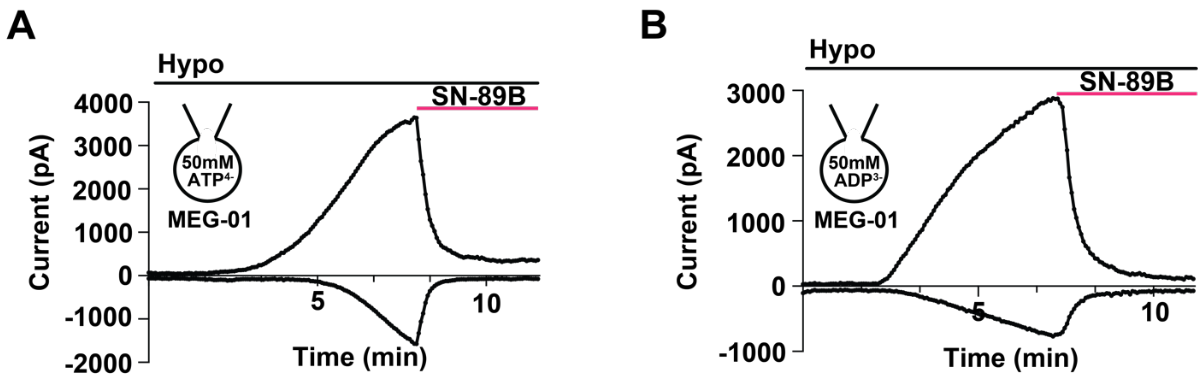


**Figure S8. LRRC8 channel inhibitor SN-89B inhibits ATP and ADP currents in MEG-01 cells. a** Current-time relationship of inward $I_{ATP}$ (ATP efflux) and outward VRAC in MEG-01 cells induced by hypotonic (210 mOsm) swelling with an intracellular ATP concentration of 50 mM, followed by application of 10 μM SN-89B. The inward component of the current represents $I_{ATP}$ generated from ATP efflux**. b** Current-time relationship of inward $I_{ADP}$ (ADP efflux) and outward VRAC in MEG-01 cells induced by hypotonic (210 mOsm) swelling with an intracellular ADP concentration of 50 mM, followed by application of 10 μM SN-89B. The inward component of the current represents $I_{ADP}$ generated from ADP efflux**.**
