## Supplementary figures and images for "LRRC8 complexes are adenosine nucleotide release channels regulating platelet activation and arterial thrombosis"

### Visual Abstract

**Visual Abstract**

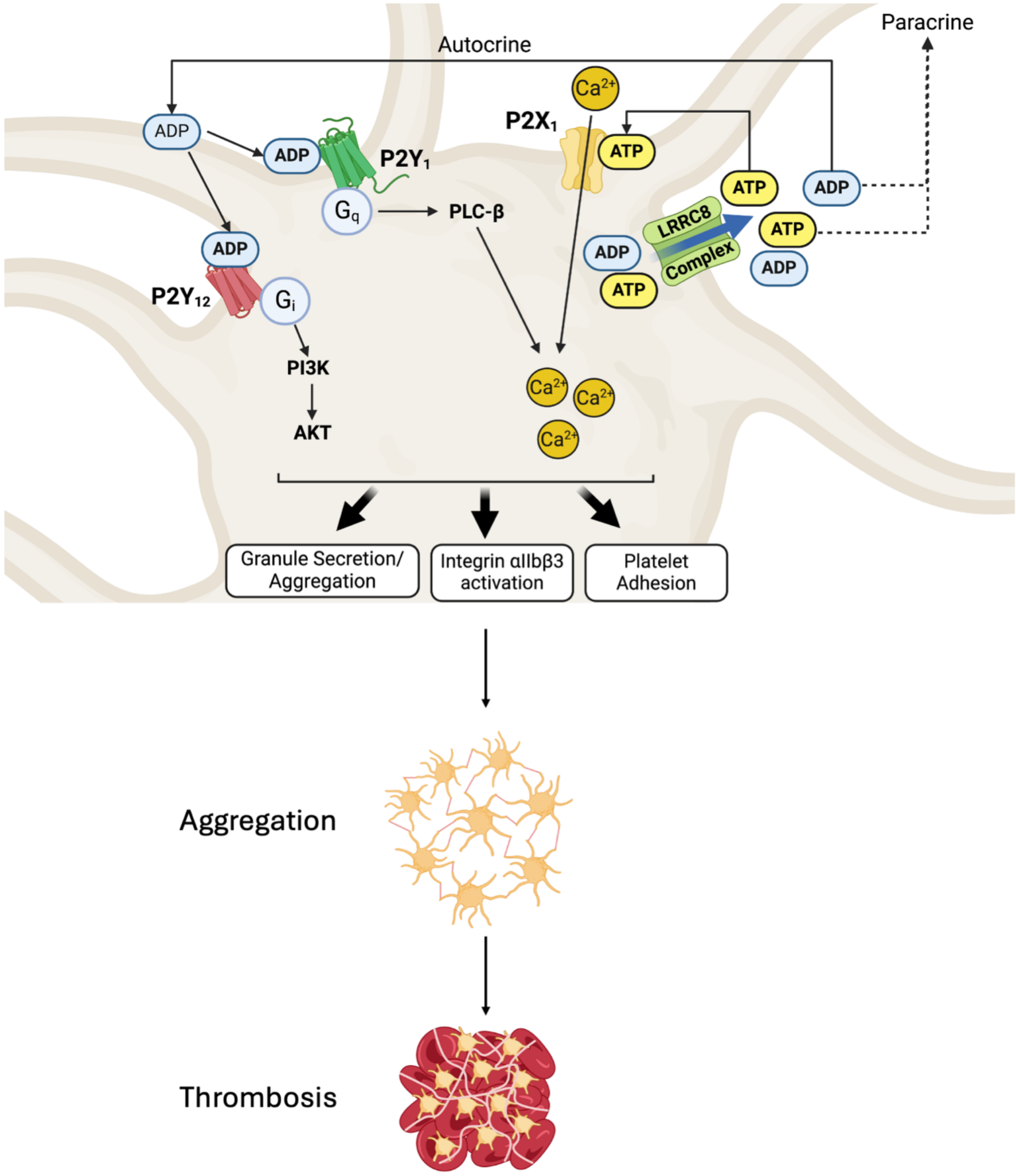
